# Transposon mutagenesis libraries reveal novel molecular requirements during CRISPR RNA-guided DNA integration

**DOI:** 10.1101/2023.01.19.524723

**Authors:** Matt W.G. Walker, Sanne E. Klompe, Dennis J. Zhang, Samuel H. Sternberg

**Author notes:** To whom correspondence should be addressed. Correspondence may also be addressed to Sanne E. Klompe. Joint Authors (Matt W.G. Walker, Sanne E. Klompe).

## Abstract

CRISPR-associated transposons (CASTs) direct DNA integration downstream of target sites using the RNA-guided DNA binding activity of nuclease-deficient CRISPR-Cas systems. Transposition relies on several key protein-protein and protein-DNA interactions, but little is known about the explicit sequence requirements governing efficient transposon DNA integration activity. Here, we exploit pooled library screening and high-throughput sequencing to reveal novel sequence determinants during transposition by the Type I-F *Vibrio cholerae* CAST system. On the donor DNA, large mutagenic libraries identified core binding sites recognized by the TnsB transposase, as well as an additional conserved region that encoded a consensus binding site for integration host factor (IHF). Remarkably, we found that VchCAST requires IHF for efficient transposition, thus revealing a novel cellular factor involved in CRISPR-associated transpososome assembly. On the target DNA, we uncovered preferred sequence motifs at the integration site that explained previously observed heterogeneity with single-base pair resolution. Finally, we exploited our library data to design modified transposon variants that enable in-frame protein tagging. Collectively, our results provide new clues about the assembly and architecture of the paired-end complex formed between TnsB and the transposon DNA, and inform the design of custom payload sequences for genome engineering applications of CAST systems.

## INTRODUCTION

Transposons are pervasive genetic elements capable of mobilizing between distinct genetic contexts using various targeting pathways, which serve as a potent force for genome evolution in all domains of life (1–3). Despite their abundance and sheer diversity, most transposons share a common feature in the presence of inverted repeat sequences that dictate the boundaries of the mobile element (4). These terminal transposon sequences are referred to as the left and right ends and typically encode one or more binding sites recognized by oligomeric transposase proteins through specific protein-DNA interactions.

DNA transposons (Class II) may be classified by their hallmark transposase domain, leading to major classes of DD(E/D) transposons, serine transposons, tyrosine transposons and Y1/Y2-transposons (5). DD(E/D)-family transposable elements encompass all known examples of “cut-and-paste” DNA transposons, which are excised from the donor site and inserted into the target site. This reaction relies on mechanistic and enzymatic symmetry, with transposase subunits executing identical chemical steps on both transposon ends that involve coordinated nucleophilic attacks during strand cleavage and joining (4). The transposase binding sites themselves, however, are not always positioned symmetrically. Whereas some transposons encode symmetrically-positioned transposase binding sites within otherwise identical left and right ends, including *Tc1/mariner-*family transposons such as *Sleeping Beauty* and Mos1 (6, 7), other elements encode asymmetrically-positioned binding sites within distinct left and right ends, including *Hermes, piggyBac*, Bacteriophage Mu, *P* elements, and Tn*7*-family transposons (8–12). These asymmetric transposon ends often facilitate strict control over integration orientation, as in the well-characterized Tn*7* transposon (8, 13), allowing one transposon end to be preferentially integrated adjacent to a target site, but the underlying mechanistic basis explaining this preference is unresolved.

Tn*7*-family transposons exhibit a modular nature in that they encode highly similar core transposition proteins — TnsA, TnsB, and TnsC — but distinct target site selection components. TnsB is a DDE-family integrase that recognizes and binds the transposon ends, catalyzes 3’ cleavage of both transferred strands at the donor site, and catalyzes the transesterification reaction at the target site. TnsA is an endonuclease protein that forms a heteromeric complex with TnsB and cleaves the 5’ end of the non-transferred strand, collectively allowing for full excision of the transposon from the donor site (14, 15). TnsC is an AAA+ ATPase protein that communicates between the transposase module and the targeting component (16, 17). Distinct Tn*7*-like transposons encode a sequence-specific DNA-binding protein as their targeting module, often of the TniQ family and referred to as TnsD, which directs transposition to a safe-harbor locus such as *glmS, comM, yhiN* and *parE* (18–23). In contrast, CRISPR-associated transposons (CASTs) use nuclease-deficient CRISPR-Cas systems to catalyze programmable, RNA-guided DNA transposition (19, 24–28).

We previously reported RNA-guided transposition activity for VchCAST, a Tn*7*-like transposon from *Vibrio cholerae* (also referred to as Tn*6677*), which mediates efficient and highly specific DNA integration in an *E. coli* heterologous host (24). VchCAST encodes a Type I-F CRISPR-Cas system that specifies integration sites through RNA-guided DNA targeting by a multi-subunit complex called Cascade (24, 29). Importantly, Cascade binds DNA in complex with an accessory transposition protein, TniQ, which ultimately recruits TnsC and the heteromeric TnsAB transposase, in complex with the donor DNA, to assemble the catalytically active transpososome (Figure 1A) (29, 30). DNA integration occurs roughly 50-bp downstream of the R-loop formed by TniQ-Cascade, but the sequence determinants underlying heterogeneous integration distances observed across distinct target sites remained enigmatic in our previous work (24, 31). Additionally, the role and requirement for multiple putative TnsB binding sites within both transposon ends is unclear, limiting efforts to further engineer VchCAST as a DNA integration tool. Like other Tn*7*-family transposons, the transposon left and right ends exhibit distinct binding site patterns, and this asymmetric arrangement may also relate to the biased orientation with which transposon insertions occur (8).

**Figure 1.**
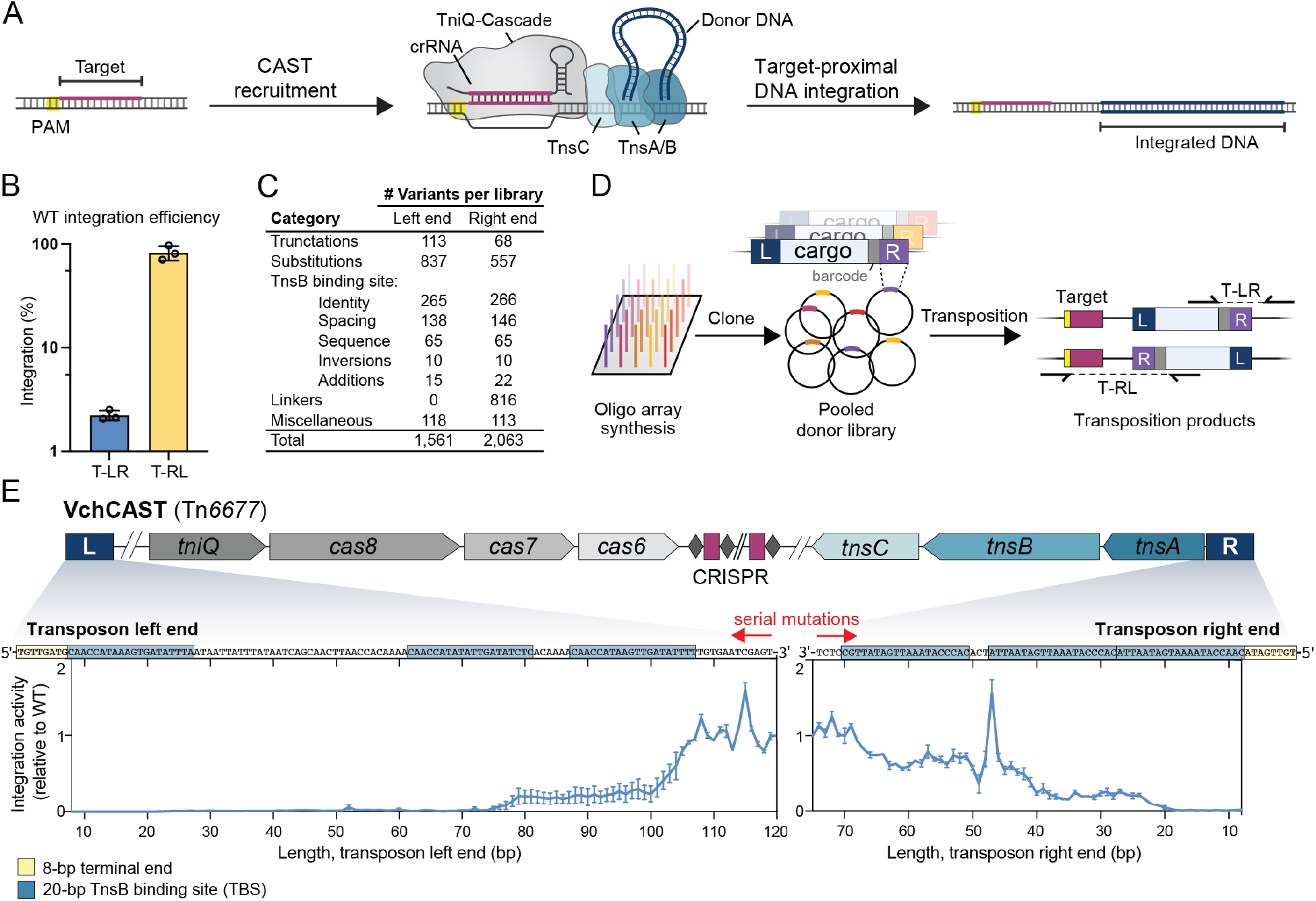
Pooled library approach to investigate transposon end mutability. (**A**) Schematic of RNA-guided transposition with VchCAST. (**B**) Integration efficiency of the WT mini-transposon in both orientations when directed to a genomic *lacZ* target site, as measured by qPCR. (**C**) Number of transposon right and left end library variants tested in each category. (**D**) Pooled library transposition approach. Library members were synthesized as single-stranded oligos and cloned into a plasmid donor library (pDonor), with 8-bp barcodes (gray) located between the transposon end and cargo used to uniquely identify each variant. The donor library was used for transposition into the *E. coli* genome, and junction amplicons were generated to determine the representation of each library member within integrated products by NGS. (**E**) Schematic of the native VchCAST system from *Vibrio cholerae* (top), and relative T-RL integration activity for library members in which the left and right ends were sequentially mutagenized beginning internally (bottom). Each point represents the average activity from two transposition experiments using the same pooled donor library.

Here, we employ library-based experiments in combination with high-throughput sequencing to investigate DNA sequence requirements during RNA-guided transposition by VchCAST. By systematically mutating both transposon ends and measuring resulting DNA integration activity, we empirically identified predicted transposase binding sites and revealed sequence preferences that mediate transposase-transposon cognate specificity. Interestingly, our results indicated that the relative positioning of each transposase binding site plays a crucial role in defining the proper architecture of the transpososome complex, with spacing patterns that correspond to the helical pitch of double-stranded DNA. These mutational data also revealed the importance of an integration host factor (IHF) binding site within the left transposon end, and subsequent genetic knockout and rescue experiments confirmed the role of IHF in stimulating transposition efficiency in *E. coli*. Finally, we uncovered new sequence preferences at the site of integration, and we exploited our mutagenesis data to rationally engineer the transposon right end to enable in-frame tagging of endogenous protein-coding genes. Collectively, this work expands our understanding of both protein and DNA sequence requirements of Tn*7*-like transposons, reveals insights into the architecture of the transpososome complex, and provides new knowledge to inform the design of custom transposon sequences for genome engineering applications.

## MATERIAL AND METHODS

### Cloning, testing, and analysis of pooled pDonor libraries

Donor plasmid (pDonor) libraries were generated by cloning transposon left or end variants into a donor plasmid, which was co-transformed with an effector plasmid (pEffector) that directed transposition into the *E. coli* genome (schematized in Figure 1D). Each transposon end variant was associated with a unique 10-bp barcode that was used to uniquely identify variants in our sequencing approach, which relied on sequencing the starting plasmid libraries (input) and integrated products from genomic DNA (output) by NGS to determine the representation of each library member before and after transposition. To sequence the output, we independently amplified integration events in the T-RL and T-LR orientations using a cargo-specific primer flanking the transposon end and a genomic primer either upstream or downstream of the integration site. We wrote custom python scripts to compare each library member’s representation in the output to its representation in the input, allowing us to calculate the relative transposition efficiency of our custom transposon end variants.

To clone the transposon donor libraries, we first generated library variants as 200-nt single stranded pooled oligos (Twist Bioscience). 1 ng of oligoarray library DNA was PCR amplified for 12 cycles in 40 μL reactions using Q5 High-Fidelity DNA Polymerase (NEB) and primers specific to the right or left end library, in order to add restriction enzyme digestion sites. Amplicons were cleaned up and eluted in 45 μL mQ H_2_O (QIAquick PCR Purification Kit). As the backbone vector, we used a plasmid encoding a 775-bp mini-transposon, delineated by 147-bp of the native transposon left end and 75-bp of the native transposon right end, on a pUC57 backbone. The backbone vector and library insert amplicons were digested (AscI and SapI for the right end library, and NcoI and NotI for the left end library) at 37 °C for 1 h, gel purified, and ligated in 20 μL reactions with T4 DNA Ligase (NEB) at 25 °C for 30 min. Ligation reactions were cleaned up and eluted in 10 μL mQ H_2_O (MinElute PCR Purification Kit), and then used to transform electrocompetent NEB 10-beta cells in five individual electroporation reactions according to the manufacturer’s protocol. After recovery (37 °C for 1 h), transformed cells were plated on large 245 mm x 245 mm bioassay plates containing LB-agar with 100 μg/mL carbenicillin. Plates were scraped to collect cells, and plasmid DNA was isolated using the QIAGEN Plasmid Midi Kit.

Transposition experiments were performed in *E. coli* BL21(DE3) cells. pEffector encoded a CRISPR array (repeat-spacer-repeat), a native *tniQ*-*cas8*-*cas7*-*cas6* operon, and a native *tnsA*-*tnsB*-*tnsC* operon, all under the control of a single T7 promoter on a pCDFDuet-1 backbone (31). 2 μL of DNA solution containing 200 ng of pDonor and pEffector in equal molar amount was used to co-transform electrocompetent cells according to the manufacturer’s protocol (Sigma-Aldrich). Four transformations were performed for each sample, and following recovery at 37 °C for 1 h, each transformation was plated on a large bioassay plate containing LB-agar with 100 μg/mL spectinomycin, 100 μg/mL carbenicillin, and 0.1 mM IPTG. Cells were grown at 37 °C for 18 h. Thousands of colonies were scraped from each plate, and genomic DNA was extracted using the Wizard Genomic DNA Purification Kit (Promega).

Next-generation sequencing (NGS) amplicons were prepared by PCR amplification using Q5 High-Fidelity DNA Polymerase (NEB). 250 ng of template DNA was amplified in 15 cycles during the PCR1 step. PCR1 samples were diluted 20-fold and amplified in 10 cycles during the PCR2 step. PCR1 primer pairs contained one pDonor backbone-specific primer and one transposon-specific primer (input library), or one genomic target-specific primer and one transposon-specific primer (output library). PCR amplicons were resolved by 2% agarose gel electrophoresis and gel-purified (QIAGEN Gel Extraction Kit). Libraries were quantified by qPCR using the NEBNext Library Quant Kit (NEB). Sequencing for both input and output libraries were performed using a NextSeq Mid or High Output Kit with 150-cycles (Illumina). Additionally, the input libraries were also sequenced using a MiSeq with 300-cycles (Illumina).

NGS data analysis was performed using custom Python scripts. Demultiplexed reads were filtered to remove reads that did not contain a perfect match to the 19-bp primer binding sequence at the 3’-terminus of the transposon end. Then, the 10-bp sequence directly downstream of the primer binding sequence was extracted, which encodes a barcode that uniquely identifies each transposon end variant. The number of reads containing each library member barcode was counted. If a read did not contain a barcode that matched a library member barcode, it was discarded. The barcode counts were summed across two NGS runs using the same PCR2 samples for the input libraries. Two biologically independent replicates were performed for the output libraries. The relative abundance of each library member was then determined by dividing the barcode count of each library member by the total number of barcode counts. The fold-change between the output and input libraries was calculated by dividing the relative abundance of each library member in the output library by its relative abundance in the input library. This fold-change was then normalized by dividing the fold-change of each library member by the average fold-change of four wildtype library members that contained identical transposon ends but unique barcodes.

One source of experimental noise in our approach came from PCR recombination (32), in which barcodes became uncoupled from their associated transposon end variants during PCR amplification. We quantified the frequency of uncoupling by performing long-read Illumina sequencing (MiSeq, 250 cycles) to sequence both the barcode and full-length transposon end, and found that not all barcodes were coupled to their correct transposon end sequence (Figure S1B). However, uncoupled reads mapped to a diverse pool of sequences, with the most abundant incorrect sequence for each library member representing only a low percentage of total reads (Figure S1C). These data therefore indicate that uncoupling events did not largely affect the ability to calculate relative integration efficiencies for each library member.

Sequence logos were generated with WebLogo 3.7.4, and the VchCAST sequence logo in Figure 2B was generated from the six predicted TnsB binding sites.

**Figure 2.**
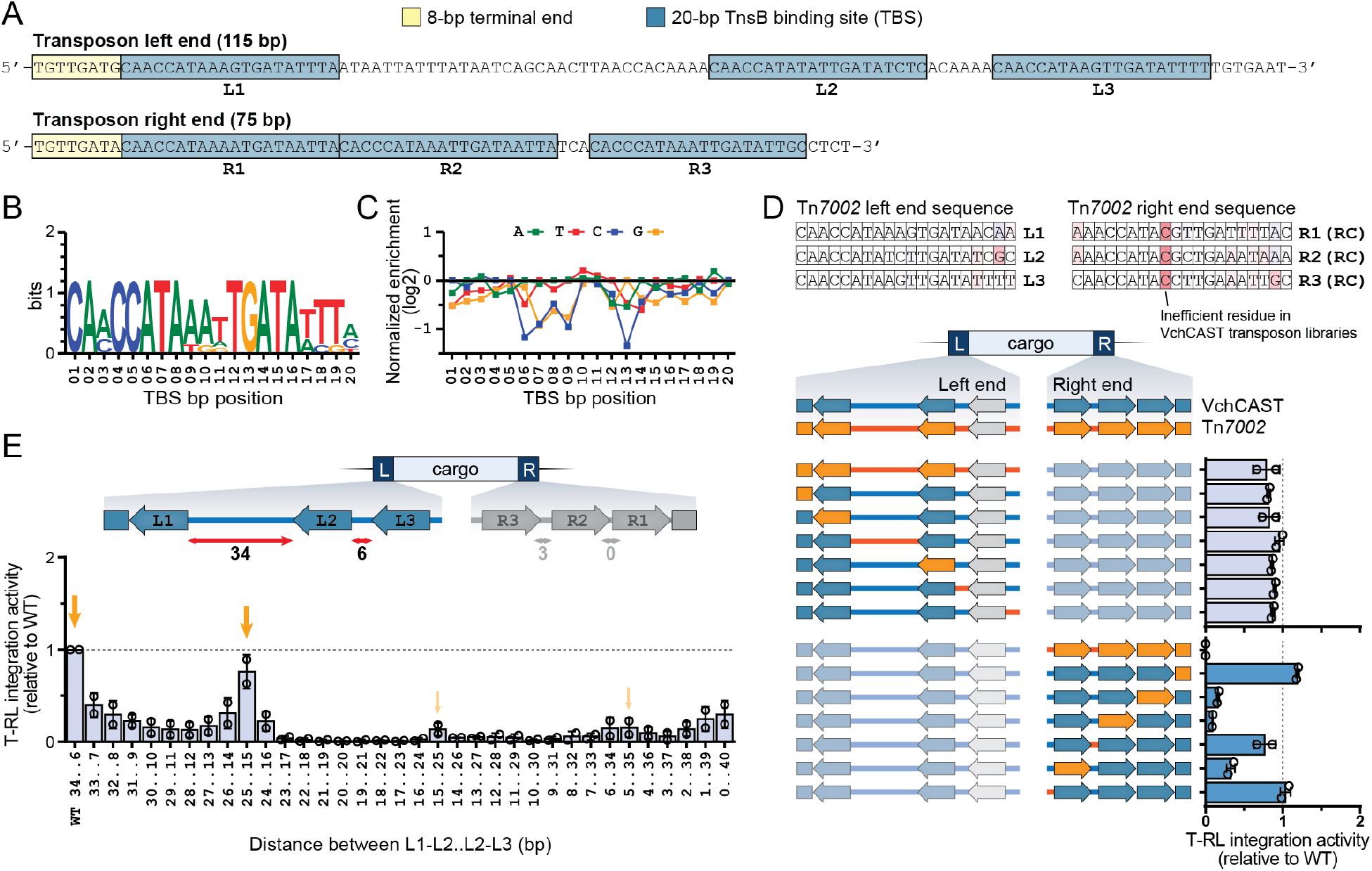
Transposase binding site (TBS) requirements for VchCAST. (**A**) Schematic representation of the VchCAST transposon end sequences. Bioinformatically predicted transposase binding site (TBS) sequences are indicated with blue boxes and labeled L1-L3 and R1-R3. The 8-bp terminal end sequences that dictate the transposon boundaries are marked with yellow boxes. (**B**) WebLogo depicting the sequence conservation of the six bioinformatically predicted TBSs. (**C**) Relative integration efficiencies (log2-transformed) for mutagenized TBS sequences averaged over all six binding sites, shown as the mean for two biological replicates. (**D**) Top: Tn*7002* transposon end sequences are colored based VchCAST transposon end library data, where red indicates a relatively inefficient residue. Bottom: relative integration efficiencies of VchCAST/Tn*7002* chimeric ends verify critical compatibility sequence requirements of TBSs. Data are shown for two biological replicates. (**E**) Relative integration efficiencies for transposon variants containing altered distances between the indicated TBSs. Orange arrows highlight the 10-bp periodic pattern of activity. Data are shown for two biological replicates.

One limitation of our experimental setup is that we could not directly compare relative integration orientation within the same NGS libraries, since integration events were amplified independently in the T-RL and T-LR orientations. Instead, we inferred approximate integration efficiencies by comparing the enrichment scores of transposon end variants to those of wildtype variants within the same library. We also note that our strategy involved separate mutagenesis of either the left end or right end, but not both simultaneously. Finally, we stress that all transposition assays with pDonor libraries were performed heterologously in *E. coli* under overexpression conditions, and thus subtleties of transposon end recognition and binding that depend on regulated TnsB expression levels may be obscured.

### Cloning, testing, and analysis of pooled pTarget libraries

pTarget libraries were designed to include an 8-bp degenerate sequence positioned 42-bp downstream of one of two potential target sites, as schematized in Figure 3B. Integration was directed to one of the two target sites flanking the degenerate sequence by a single plasmid (pSPIN) encoding both the donor molecule and transposition machinery under the control of a T7 promoter, on a pCDF backbone [described in (33)]. To generate insert DNA for cloning the pTarget libraries, two partially overlapping oligos (oSL2241 and oSL2245, Table S2) were annealed by heating to 95 °C for 2 min and then cooling to room temperature. Annealed DNA was treated with DNA Polymerase I, Large (Klenow) Fragment (NEB) in 40 μL reactions and incubated at 37 °C for 30 min, then gel-purified (QIAGEN Gel Extraction Kit). Double-stranded insert DNA and vector backbone was digested with BamHI and AvrII (37 °C, 1 h); the digested insert was cleaned-up (MinElute PCR Purification Kit) and the digested backbone was gel-purified. Backbone and insert were ligated with T4 DNA Ligase (NEB), and ligation reactions were used to transform electrocompetent NEB 10-beta cells in four individual electroporation reactions according to the manufacturer’s protocol. After recovery (37 °C for 1 h), cells were plated on large bioassay plates containing LB-agar with 50 μg/mL kanamycin. Thousands of colonies were scraped from each plate, and plasmid DNA was isolated using the QIAGEN Plasmid Midi Kit. Plasmid DNA was further purified by mixing with Mag-Bind TotalPure NGS Beads (Omega) at a vol:vol ratio of 0.60 x and extracting the supernatant to remove contaminating fragments smaller than ∼450 bp.

**Figure 3.**
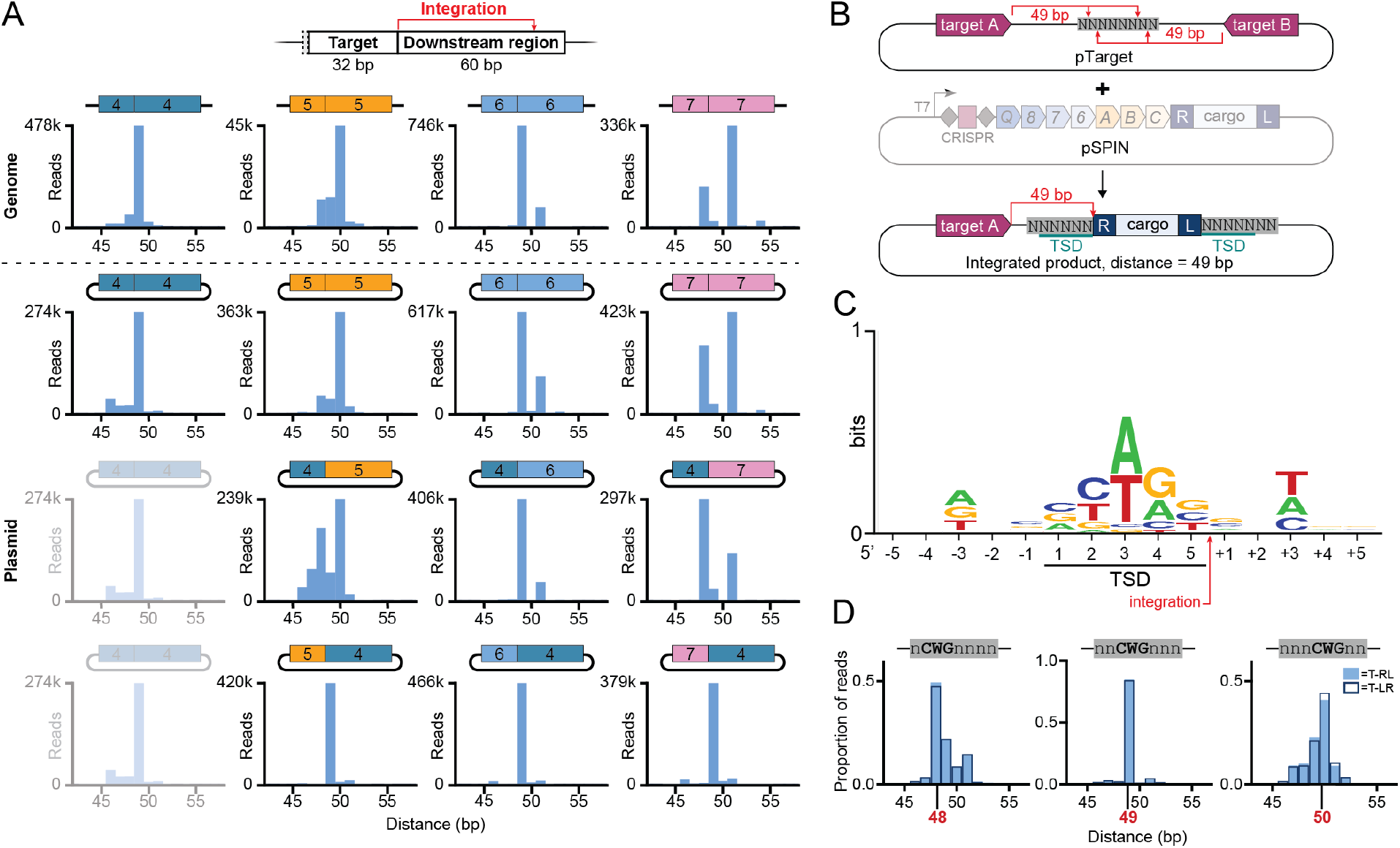
Transposase sequence preferences influence integration site patterns. (**A**) VchCAST exhibits target-specific heterogeneity in the distance (d) between the target site and integration site, which could result from sequence preferences within the downstream region (top). distances Deep sequencing revealed biases in integration site preference, with integration patterns shown for four target sites (4–7) located in the *lac* operon of the *E. coli* BL21(DE3) genome (top row) or encoded on a separate target plasmid (second row). Chimeric target plasmids that either maintain the 32-bp target site (third row) or 60-bp downstream region (bottom row) of target 4 were also tested. These data reveal that sequence identity of the downstream region (including the integration site), but not the target site, governs the observed in integration distance distribution. (**B**) Schematic of integration site library experiment, in which integration was directed into an 8-bp degenerate sequence encoded on a target plasmid (pTarget). (**C**) Sequence logo of preferred integration site, generated by selecting nucleotides from the top 5000 enriched sequences across all integration positions in each library, with a minimum threshold of four-fold enrichment in the integrated products compared to the input. (**D**) The preferred 5’-CWG-3’ motif in the center of the TSD is predictive of integration site distribution, as the displacement of this motif within the degenerate sequence shifts the preferred integration site distance, indicated by the red number.

2 μL of DNA solution containing 200 ng of pTarget and pSPIN at equal mass amounts were used to co-transform electrocompetent *E. coli* BL21(DE3) cells according to the manufacturer’s protocol (Sigma-Aldrich). Three transformations were performed and plated on large bioassay plates containing LB-agar with 100 μg/mL spectinomycin and 50 μg/mL kanamycin. Thousands of colonies were scraped from each plate, and plasmid DNA was isolated using the QIAGEN Plasmid Midi Kit.

Integration into pTarget yielded a larger plasmid than the starting input plasmid. To isolate the larger plasmid, we performed a digestion step that facilitated resolution of the integrated and unintegrated bands on an agarose gel, for extraction of the larger integrated plasmid. We performed this digestion step on both input and output libraries, digesting with NcoI-HF (37 °C for 1 h) and running them on a 0.7% agarose gel. The products were gel-purified (QIAGEN Gel Extraction Kit) and eluted in 15 μL EB in a MinElute Column (QIAGEN). 6.5 μL of cleaned-up DNA was used in each PCR1 amplification with Q5 High-Fidelity DNA Polymerase (NEB) for 15 cycles. PCR1 samples were diluted 20-fold and amplified in 10 cycles for PCR2. PCR1 primer pairs contained pTarget backbone-specific primers flanking a 45-bp region encompassing the degenerate sequence. Sequencing was performed with a paired-end run using a NextSeq High Output Kit with 150-cycles (Illumina).

NGS data analysis was performed using a custom Python script. Demultiplexed reads were filtered to remove reads that did not contain a perfect match to the 34- to 35-bp sequence upstream of the degenerate sequence for any i5-reads, or to the 45- to 46-bp sequence for any i7-reads. (35-bp and 46-bp was used for reads that were amplified from primers containing an additional nucleotide, which were used in PCR1 to generate cluster diversity during sequencing). For all reads that passed filtering, the 8-bp degenerate sequence was extracted and counted. The integration distance was determined in the output libraries by examining the i5 read sequence at an integration distance of 43-bp to 56-bp downstream of each target for the presence of the transposon right or left end sequence (20-nt of each end). The degenerate sequence was then extracted from either or both of the i5 and i7 reads, depending on the integration position. The degenerate sequence counts were summed across the two primer pairs. The relative abundance was determined by dividing the degenerate sequence count by the total number of degenerate sequence counts. Finally, the fold-change between the output and input libraries was calculated by dividing the relative abundance of each degenerate sequence at each integration position in the output library by its relative abundance in the input library, and then log2-transformed.

Sequence logos were generated with WebLogo 3.7.4. The preferred integration site logos in Figure S3A were generated from all degenerate sequences that were enriched four-fold in the integrated products compared to the input. The overall preferred integration site logos in Figure 3C and Figure S3D were generated by first applying the minimum threshold of four-fold enrichment in the integrated products compared to the input, and then selecting nucleotides from the top 5,000 enriched sequences across all integration positions. We selected nucleotides from the top 5,000 sequences from each library, yielding a total of 10,000 nucleotides at each position.

### Endogenous gene tagging experiments

All VchCAST constructs were subcloned from pEffector and pDonor as described previously, using a combination of inverse (around-the-horn) PCR, Gibson assembly, restriction digestion-ligation, and ligation of hybridized oligonucleotides (24, 31). pEffector encodes a CRISPR array (repeat-spacer-repeat), a native *tniQ*-*cas8*-*cas7*-*cas6* operon, and a native *tnsA*-*tnsB*-*tnsC* operon, all under the control of a single T7 promoter on a pCDFDuet-1 backbone (31). Donor plasmids (pDonor) were designed to encode a mini-transposon (mini-Tn) with a wild-type 147-bp transposon left end and 57-bp linker-coding right end variant, on a pUC19 backbone. For endogenous gene tagging experiments, superfolder GFP (sfGFP) lacking a ribosome binding site (rbs) and start codon was cloned into the mini-Tn cargo region, and the mini-Tn was further cloned into a temperature-sensitive pSIM6 backbone.

Linker functionality constructs were designed to encode sfGFP with an extended 32-amino acid (aa) loop region between the 10^th^ and 11^th^ β-strands, under the control of a single T7 promoter, as described by Feng and colleagues (34). Linker variants encoding 18-19 aa were subcloned into the 32-aa loop region as follows. An entry vector was generated on a pCOLADuet-1 (pCOLA) vector harboring sfGFP, such that the 11th β-strand (GFP11) was replaced by the aforementioned extended 32-aa loop (34). Fragments encoding transposon right end linker variants and GFP11 were then amplified by conventional PCR and inserted into the extended loop region of the entry vector downstream of β-strands 1–10 (GFP1-10), such that total length of the loop remained constant at 32 aa.

To perform linker functionality assays, chemically competent *E. coli* BL21(DE3) cells were co-transformed with T7-controlled sfGFP linker functionality constructs (pCOLA) and an equal mass amount of empty pUC19 vector. Negative control transformants harbored either unfused sfGFP1-10 and sfGFP11 fragments on separate pCOLA and pUC19 backbones, respectively, or isolated sfGFP fragments. Transformed cells were plated on LB-agar plates with antibiotic selection (100 μg/mL carbenicillin, 50 μg/mL kanamycin), and single colonies were used to inoculate 200 μL of LB medium (100 μg/mL carbenicillin, 50 μg/mL kanamycin, 0.1 mM IPTG) in a 96-well optical-bottom plate. The optical density at 600 nm (OD600) was measured every 10 min, in parallel with the fluorescence signal for sfGFP, using a Synergy Neo2 microplate reader (Biotek) while shaking at 37 °C for 15 h. To derive normalized fluorescence intensities (NFI), all measured fluorescence intensities were divided by their corresponding OD600 values across all time points. A single representative NFI value was calculated per well by averaging all NFI values per well corresponding to OD600 values between 0.20 and 0.30, inclusive.

Transposition experiments were performed by transforming chemically competent *E. coli* BL21(DE3) cells harboring pEffector plasmids with pDonor plasmids by heat shock at 42 °C for 30 sec, followed by recovery in fresh LB medium. Recovery was performed at 30 °C for 1.5 h for temperature-sensitive pDonor plasmids, and 37 °C for 1 h for all other pDonor plasmids. Transformants were isolated on LB-agar plates containing the proper antibiotics and inducer (100 μg/mL carbenicillin, 50 μg/mL spectinomycin, 0.1 mM IPTG). After 43 h growth at 30 °C for temperature-sensitive pDonor plasmids, and 18 h growth at 37 °C for all other pDonor plasmids, samples were prepared for downstream qPCR analysis of integration efficiency or colony PCR identification of integration events.

For qPCR quantification, colonies were scraped from plates and resuspended in LB medium, and cell lysates were prepared for qPCR as described by Klompe and colleagues (24). Pairs of transposon- and target DNA-specific primers were designed to amplify fragments from integrated transposition products at the expected loci in either of two possible orientations. In parallel, a separate pair of genome-specific primers was designed to amplify an *E. coli* reference gene (*rssA*) for normalization purposes. qPCR reactions (10 μL) contained 5 μL of SsoAdvanced Universal SYBR Green Supermix (BioRad), 1 μL H2O, 2 μL of 2.5 μM primers, and 2 μL of hundredfold-diluted cell lysate and were prepared following transposition experiments as described above. Reactions were prepared in 384-well clear/white PCR plates (BioRad), and measurements were obtained in a CFX384 Real-Time PCR Detection System (BioRad). The following thermal cycling parameters were used: polymerase activation and DNA denaturation (98 °C for 3 min), and 35 cycles of amplification (98 °C for 10 s, 60 °C for 30 s). Each biological sample was analyzed in three parallel reactions: one reaction contained a primer pair for the *E. coli* reference gene, a second reaction contained a primer pair for one integration orientation, and a third reaction contained a primer pair for the other integration orientation. Transposition efficiency was calculated for each orientation as 2ΔCq, in which ΔCq is the Cq difference between the experimental and control reactions. Total transposition efficiency for a given experiment was calculated by summing transposition efficiencies across both orientations. All measurements presented were determined from three independent biological replicates.

For colony PCR identification of integration events, colonies were scraped from plates after transposition assays, resuspended in fresh LB medium, and re-streaked on LB-agar plates with the appropriate antibiotics and without IPTG inducer. To generate lysates, individual colonies were each transferred to 10 μL of H2O, followed by incubation at 95 °C for 2 min and centrifugation at 4,000 g for 5 min to pellet cell debris. Pairs of transposon- and target DNA-specific primers were designed to amplify fragments from integrated transposition products in the expected locus and orientation. In parallel, a separate pair of genome-specific primers was designed to amplify an *E. coli* reference gene (*rssA*) and determine whether the crude lysates were sufficiently dilute to allow successful amplification of the integrated transposition product. Transposition-less negative control samples were always analyzed in parallel with experimental samples to identify mispriming products that could result from the pDonor-containing crude lysates. PCR reactions (15 μL) contained 7.5 μL of 2X OneTaq 2X Master Mix with Standard Buffer (NEB), 5.9 μL H2O, 0.6 μL of 10 μM primers, and 1 μL of undiluted cell lysate as described above. PCR amplicons were resolved by 1% agarose gel electrophoresis and visualized by staining with SYBR Safe (Thermo Scientific). To verify in-frame integration events, amplicons of the expected length were excised after gel electrophoresis, isolated by the Gel Extraction Kit (Qiagen), and sent for Sanger sequencing (GENEWIZ).

Fluorescence microscopy experiments were performed as follows. A pEffector plasmid was designed to C-terminally tag the native *E. coli msrB* gene by integrating a mini-Tn encoding a linker variant (ORF2a) and sfGFP cargo in-frame with the coding sequence, thereby interrupting the endogenous stop codon. Transposition experiments were performed as described above by transforming chemically competent *E. coli* BL21(DE3) cells harboring pEffector plasmids with temperature-sensitive pDonor plasmids. Colonies were then scraped and resuspended in fresh LB medium. Resuspensions were diluted and re-streaked on double antibiotic LB-agar plates lacking IPTG (100 μg/mL carbenicillin, 50 μg/mL spectinomycin). After overnight growth on solid medium at 37 °C, individual colonies were used to inoculate liquid cultures (50 μg/mL spectinomycin) for overnight heat-curing at 37 °C, followed by replica plating on single and double antibiotic plates to isolate heat-cured samples. In tandem, colony PCR and Sanger sequencing (GENEWIZ) were performed to identify colonies with in-frame transposition products as described above. In preparation for fluorescence microscopy, Sanger-verified samples were inoculated in overnight 37 °C liquid cultures. On the day of imaging, 500 μL of saturated overnight cultures were transferred to 5 ml of fresh LB medium with the appropriate antibiotics. Aliquots of the newly inoculated cultures were removed around the stationary or mid-log phases and immobilized in glass slides coated with partially dehydrated aqueous 1% agarose-TAE pads. Immediately after immobilization, fluorescent microscopy was performed with a Nikon ECLIPSE 80i microscope using an oil immersion x100 objective lens, which was equipped with a Spot CCD camera and SpotAdvance software. All images were processed in ImageJ by normalizing background fluorescence.

### Generating and testing *E. coli* knockout mutants

*E. coli* genomic knockouts of *ihfA, ihfB, ycbG, hupA, hupB, hns*, and *fis* were generated using Lambda Red recombineering, as previously described (35). Knockouts were designed to replace of each gene with a kanamycin resistance cassette, which was PCR amplified with Q5 High-Fidelity DNA Polymerase (NEB) using primers that contained 50-nt homology arms to knockout gene locus. PCR amplicons were resolved on a 1% agarose gel and gel-purified, eluting with 40 μL MQ (QIAGEN Gel Extraction Kit). Electrocompetent *E. coli* BL21(DE3) cells were prepared containing a temperature-sensitive plasmid that encodes the Lambda Red machinery under the control of a temperature-sensitive promoter (pSIM6). Protein expression from the temperature-sensitive promoter was induced by incubating cells at 42 °C for 25 min immediately prior to electrocompetent cell preparation. 300-600 ng of each insert was used to transform cells via electroporation (2 kV, 200 Ω, 25 μF), and cells were recovered overnight at 30 °C by shaking in 3 mL of SOC media. After recovery, 250 μL of culture was spread on 100 mm standard plates (LB-agar with 50 μg/mL kanamycin) and grown overnight at 30 °C. Kanamycin-resistant colonies were picked, and the genomic knock-in was confirmed by PCR amplification and Sanger sequencing using primer pairs flanking the knock-in locus.

VchCAST transposition experiments in *E. coli* knockout strains were performed by first preparing chemically competent WT and mutant cells and then transforming these strains with a single plasmid (pSPIN), which encodes the donor molecule and the native transposition machinery under the control of a T7 promoter and a crRNA targeting the *lacZ* genomic locus, on a pCDF backbone. After transformation by heat shock, cells were plated onto LB-agar with 100 μg/mL spectinomycin and 0.1 mM IPTG to induce protein expression, and incubated at 37 °C for 18 h. Hundreds of colonies were scraped from each plate, and integration efficiencies were quantified by the same qPCR assay described for the endogenous gene tagging experiments. Transposition experiments for other Type I-F homologs were performed as in the VchCAST experiments, except that the concentration of IPTG was reduced to 0.01 mM to mitigate toxicity.

Experiments that tested protein expression conditions in WT and ΔIHF cells were performed as described in the VchCAST transposition experiments. Promoters were varied from constitutive promoters (J23119, J23101) to inducible promoters (T7), for which different concentrations of IPTG were also tested.

For the complementation experiments, cells were co-transformed with pSPIN and a rescue plasmid (pRescue) that encoded both *E. coli ihfA* and *ihfB* under the control of separate T7 promoters on a pACYC backbone, and plated onto LB-agar with 100 μg/mL spectinomycin, 25 μg/mL chloramphenicol, and 0.1 mM IPTG to induce protein expression. Cells were incubated at 37 °C for 18 h, before colonies were scraped from each plate and integration efficiencies in both orientations were measured by qPCR.

To test DNA donor molecules with symmetric transposon ends, we cloned mutant pDonor encoding two right or two left transposon ends, and measured integration efficiency by co-transforming pDonor with pEffector under the control of a T7 promoter on a pCDF backbone. Cells were plated onto LB-agar with 100 μg/mL spectinomycin, 100 μg/mL carbenicillin, and 0.1 mM IPTG and incubated at 37 °C for 18 h, before colonies were scraped from each plate and integration efficiencies in both orientations were measured by qPCR.

### EcoTn7 transposition experiments and NGS analysis

To measure the integration efficiencies and distance distributions of EcoTn*7* in WT and *E. coli* mutant cells, we cloned genomic primer binding sites into the mini-Tn cargo of a single plasmid for Tn*7* transposition, which encoded a native *tnsA-tnsB-tnsC-tnsD* operon under the control of a constitutive pJ23119 promoter, on a pCDF backbone. The genomic primer binding sites were cloned adjacent to the transposon left and right ends such that the NGS amplicon length would be the same for unintegrated products and integrated products in either orientation (schematized in Figure S7A). To quantify integration efficiencies using qPCR, we used primer pairs designed to amplify integrated products in both orientations, with one primer adjacent to the right transposon end a second primer either upstream or downstream of the integration site.

To quantify integration efficiencies by NGS, we amplified genomic DNA using a single primer pair with one primer complementary to the genomic primer binding site and the second primer complementary to the 3’-end of the *glmS* locus. Genomic DNA was extracted using the Wizard Genomic DNA Purification Kit (Promega). 250 ng of genomic was used in each PCR1 amplification with Q5 High-Fidelity DNA Polymerase (NEB) for 15 cycles. PCR1 samples were diluted 20-fold and amplified in 10 cycles for PCR2. Sequencing was performed with a paired-end run using a NextSeq High Output Kit with 150-cycles (Illumina).

NGS data analysis was performed using a custom Python script. Demultiplexed reads were filtered to remove reads that did not contain a perfect match to the first 65-bp of expected sequence resulting from either non-integrated genomic products or from integration events spanning 0-bp to 30-bp downstream of the *glmS* locus, and then counted the number of reads matching each of these possible products.

## RESULTS

### A pooled library approach to investigate transposon end sequence requirements

We set out to systematically mutagenize the transposon left and right end sequences of *V. cholerae* Tn*6677* using large pooled oligoarray libraries, building off our previous study of the VchCAST system (24). Starting with a minimal pDonor design that directed efficient genomic integration in both of two possible orientations (Figure 1B), we designed thousands of variants of the left (L) and right (R) end sequences, including truncations, base-pair substitutions, and transposase binding site modifications (Figure 1C and Table S3 and S4). We assigned each variant a unique 8-bp barcode located between the mutagenized transposon end and the cargo, obviating the requirement to sequence across the entire transposon end to identify each variant. Each library also included four wildtype (WT) variants associated with unique barcodes, which we used to approximate the relative integration efficiency of each mutagenized library member. Libraries were then synthesized as single-stranded oligos, cloned into a mini-transposon donor (pDonor), and carefully characterized using next-generation sequencing (NGS), which demonstrated that all members were represented in the input sample for both transposon left and right end libraries (Figure S1A–D).

We performed transposition experiments by transforming *E. coli* BL21(DE3) cells expressing the transposition machinery with pDonor encoding either the left end or right end library, amplifying successful genomic integration products in both orientations via junction PCR (Figure 1D), and subjecting PCR products to NGS analysis. An enrichment score was then calculated for each variant, revealing a wide range of integration efficiencies, with most library members exhibiting diminished integration relative to the four WT samples (Figure S1D). Finally, we used enrichment scores of the WT library members for normalization, yielding a score for each variant that represented its relative activity. To validate our approach, we performed two biological replicates for each library transposition experiment and found strong concordance between both datasets, especially in the dominant T-RL integration orientation (Figure S1E). Importantly, we also rigorously determined the background level of library member–barcode uncoupling, given the high degree of sequence similarity between library members, which established contributors of experimental noise in our datasets (Figure S1B-C and Methods).

The strength of the pooled-library approach is apparent by examining the effect of one category of variations, in which we sequentially mutated the transposon end sequences starting 120-nt into the transposon end, effectively creating end truncations, albeit without a change in overall mini-transposon size (Figure 1E). These results revealed the minimal transposon end sequence length: in the left end, ∼105 bp were required for efficient integration, corresponding to all three predicted transposase (TnsB) binding sites, whereas in the right end, only ∼50 bp were required, corresponding to the first two transposase binding sites. These findings are consistent with previous literature and adds single-bp resolution to the minimal transposon end sequences needed for efficient integration (24).

### Transposase activity depends on specific sequence requirements

TnsB is integral to the mobilization of Tn*7*-like transposons, in that it catalyzes the excision and integration chemistry while also conferring sequence specificity for the transposon ends through recognition of repetitive sequence elements known as TnsB binding sites (TBSs) (8, 15, 36). Sequence analysis of the native VchCAST ends revealed three conserved TBSs in both the left and right ends (Figure 2A,B and Figure S2A) (24), and we verified these sequence requirements by examining a mutational panel at single-bp resolution (Figure 2C and Figure S2B). This dataset revealed that individual TBS point mutations can affect efficiency, particularly for positions 1, 6-9, and 12-14, but are not critical for integration. This more lenient sequence requirement is in line with recently published cryo-EM structures of DNA-bound TnsB from Tn*7* and Type V-K CAST systems, which revealed that many protein-DNA interactions occur with the phosphodiester backbone rather than specific nucleobases (37–39).

Experiments with *E. coli* Tn*7* showed that the internal TBSs are occupied before the more terminal sites (8). Even though the six TBSs of VchCAST differ by only a couple bases, we wondered if these differences might be biologically important, by enforcing a specific assembly pathway. To test this hypothesis, we tested all possible combinations of TBSs for the left and right ends, which we defined as L1–L3 and R1–R3 (Figure S2C). For both VchCAST ends, site 1 displayed the greatest TBS preference and preferred the L1/L3/R1 sequence, whereas site 2 preferred L1/R1/R2 and site 3 exhibited the least TBS preference but favored L3. We observed a preference for R1 in the first position on the left end, and a preference for L1 in the first position on the right end, suggesting that transposition might be favored when the terminal end sequences are identical (whether based on equal affinity or otherwise).

Apart from regulating transposition frequency, TBS sequence identity could also explain the propensity of a given CAST system to cross-react with related transposon substrates (18). We previously showed that VchCAST could efficiently mobilize mini-transposon substrates from three homologous CAST systems, but not Tn*7002*. To determine which Tn*7002* sequences were incompatible with mobilization by VchCAST machinery, we designed chimeric transposon ends that contain parts of both the VchCAST and Tn*7002* transposon ends (Figure 2D). The data revealed that chimeric left ends allowed for near WT integration efficiencies whereas chimeric right ends drastically decreased integration efficiency, likely due to the deleterious presence of a cytidine at position 9 of R1–R3 (Figure 2D). These data thus demonstrate that TBS sequence identity imparts specific constraints on the substrate recognition of a transposase for its cognate transposon DNA.

Finally, we sought to investigate the conserved positioning of TBSs within the transposon ends, after hypothesizing that the specific distance between TBSs might facilitate proper assembly of transposase subunits within a paired-end-complex (PEC) (18). After testing a mutagenic panel in which the length between TBSs was systematically varied (Figure 2E and Figure S2D), we found that even single-bp perturbations caused drastic changes in integration efficiency. Additionally, we detected an intriguing pattern of increasing and decreasing integration efficiencies at roughly 10-bp intervals, suggesting that the three-dimensional positioning of transposase proteins on helical DNA is important for transposition.

Together, these data highlight the impact of TBS mutations and TBS sequence positioning on transposition, and provide clues about how TBSs may have evolved to direct efficient assembly of synaptic paired-end complexes.

### Transposase sequence preferences influence integration site patterns

In our previous work, we showed that VchCAST integration patterns differed in subtle but reproducible ways between distinct genomic target sites (24, 31). Since integration is the result of both RNA-guided DNA targeting and transposase-mediated DNA integration, we were curious to investigate which DNA sequences and protein machineries were responsible for the heterogeneity in integration products. We first compared integration site patterns for four endogenous *E. coli* target sequences, designated 4–7, either at their native genomic location or on an ectopic target plasmid by deep sequencing (Figure 3A). Integration site patterns were notably distinct between the four targets but were highly consistent between genomic and plasmid contexts, suggesting that these patterns are dependent on local sequence alone and independent of other factors such as DNA replication or local transcription. Next, to disentangle contributions of the 32-bp target sequence (complementary to crRNA guide) from the downstream region including the integration site, we tested target plasmids that contained chimeras of the four target regions (Figure 3A). Remarkably, integration patterns for these chimeric substrates closely mirrored the patterns observed for the non-chimeric substrates when the ‘downstream region’ was kept constant, clearly indicating that the 32-bp target sequence does not modulate selection of the integration site.

We hypothesized that, like other transposases, TnsB might exhibit local sequence preferences immediately at the site of DNA insertion, and that these preferences could explain the observed heterogeneity in integration site patterns (40). To test this possibility, we generated a target plasmid (pTarget) library encoding two target sequences flanking an 8-bp degenerate sequence, such that integration events directed by a crRNA matching either target would lead to insertion directly into the degenerate 8-mer sequence (Figure 3B). We sequenced the target plasmids before and after transposition and compared the representation of integration site sequences to determine which sequences were enriched after transposition. These analyses revealed striking nucleotide preferences at conserved positions relative to the integration site (Figure 3C and Figure S3A). Specifically, there were clear biases for a YWR motif within the central three nucleotides of the target-site duplication (TSD), as well as a preference for D (A, T, or G) and H (A, T, or C) at the –3 and +3 positions relative to the TSD, respectively. Similar TSD preferences were previously observed for the Type V-K ShCAST system (25), suggesting that they may be broadly applicable to TnsB-family transposases.

To further explore the deterministic role of the preferred motif within the TSD, we plotted the distribution of reads containing a central 5’-CWG-3’ motif at different positions within the degenerate sequence. We focused on this motif because it favored a more unimodal distribution for the integration site by avoiding a centrally-preferred A or T nucleotide flanking the W. We found that this motif was indeed predictive of the preferred integration site distance that was sampled by VchCAST (Figure 3D). We extended this observation by plotting the distribution of reads containing multiple 5’-CWG-3’ motifs within the integration site and found that two copies of this preferred motif within the integration site conferred a bimodal distribution, wherein there were not one but two preferred integration sites within the degenerate sequence (Figure S3B). Finally, we leveraged our library data to predict the integration site distribution of previously targeted locations (24) and found that we could explain their differences at single-bp resolution (Figure S3E).

Both of the two distinct crRNAs and corresponding target sites on pTarget yielded consistent sequence preferences for both the TSD and +/- 3-bp positions (Figure S3A), but we were surprised to find that the preferred integration distance was shifted by 1 bp when comparing the two (Figure S3C). We suspected that this difference could be due to sequences preferences at the +/-3-bp position that fell outside the degenerate sequence, and indeed, when we examined the sequences flanking the 8-mer library, we found that the downstream target (target B) contained a disfavored nucleotide in the -3-bp position for insertions that would occur with the 49-bp distance (Figure S3D). Interestingly, the role for these positions in modulating transposition behavior is well-substantiated by a recent structure of TnsB from *Scytonema hofmanni* Type V-K CAST bound to strand-transfer intermediates (41), which showed residue K290 of both terminal TnsB protomers contacting the +/- 3-bp position of the target site.

### Role of boundary sequences and right end internal features on DNA integration

We next focused our attention on additional sequence features at the outermost edges of mini-transposon substrates. VchCAST and many other Tn*7*-like transposons encode an 8-bp terminal end immediately adjacent to the first transposase binding site, with the terminal TG dinucleotide highly conserved among a broad spectrum of transposons including IS3, Tn*7*, Mu and even retrotransposons (43–46). Integration data with library variants that featured mutations within these terminal residues revealed that positions 1–3, but not 4–8, were critical for efficient transposition (Figure S4B). This result is consistent with the DNA-bound cryo-EM structure of TnsB from a Type V-K CAST system, in which base-specific interactions were observed for the terminal TG dinucleotide (37), and with experiments indicating that these terminal dinucleotides are important for the formation of a stable Mu transpososome complex (44, 47). Sequences beyond the terminal TG are also acted upon during excision of Tn*7*-like transposons, since the endonuclease TnsA cleaves the 5’ ends of the donor DNA 3 bp outside the transposon end boundaries (42). This observation suggested the possibility that the sequence context of the transposon donor itself might play a role in efficient transposition. However, library variants with mutations in the 5-bp sequence flanking the mini-transposon were integrated with equivalent efficiencies (Figure S4A), indicating that transposition machinery does not exhibit sequence specificity within this region.

To investigate whether the spacing between the terminal TG dinucleotide and the first TBS mattered, we tested variants that modulated the distance between the 8-bp terminal end and TBS1 (Figure S4C). Adding a single base pair in either the left or right end still allowed for efficient transposition, whereas transposition was completely ablated with the removal of 1 bp or addition of 2 bp, indicating tight control over this spacing. Interestingly, larger bp additions or deletions between the TG dinucleotide and first TBS were in some cases also permitted, but always with a concomitant shift in the transposon boundary that was actually mobilized and integrated at the target site (Figure S4C); in all cases, transposition still required a terminal TG. These data therefore suggest that the critical feature within the terminal end sequence is the TG dinucleotide, and that the ∼8-bp spacing between this dinucleotide and the first TBS is a critical constraint for efficient transposition.

We also further investigated the importance of a palindromic sequence found 97–107 bp from the transposon right end boundary. Previous work suggested that this sequence might affect integration orientation, possibly by promoting transcription of the *tnsABC* operon, which would be consistent with empirical expression data and the AT-richness of the transposon end (48). To test this possibility, we mutated the palindromic sequence and found that variants with this sequence shifted the orientation preference towards T-LR, with just one arm of the palindrome (P_B_) being sufficient to shift the orientation bias (Figure S4D-E). We also included bona fide constitutive promoters in place of the palindromic sequence and found that promoters directing transcription inwards (towards the cargo) did not impact integration orientation, whereas promoters directed outwards (across the right end) shifted the orientation preference towards T-LR, perhaps by antagonizing stable assembly of TnsB selectively at the right end (Figure S4F). These data highlight the role of this right end sequence region on integration orientation, which should be considered when designing custom cargo sequences.

### Endogenous protein tagging with rationally engineered right ends

The left and right end sequences are critical for transposon DNA recognition and excision/integration, and transposition products therefore necessarily include these sequences as ‘scars’ at the site of insertion. We sought to exploit this feature and use our new knowledge of the mutability of the transposon ends to convert these scars into functional sequences that encode amino acid linkers for downstream protein tagging applications. We focused on the shorter right end, starting with a minimal 57-bp sequence, and observed that stop codons were present in all three possible open reading frames (ORF) for the WT sequence (Figure 4A) (24). When we tested a library of rationally designed right end variants that replaced stop codons and codons encoding bulky and/or charged amino acids, we identified numerous candidates for each possible ORF that maintained near-wild-type integration efficiency (Figure S5A). After validating library data by testing individual linker variants for genomic integration in *E. coli* (Figure 4B), we next set up a fluorescence-based assay to test for functionality of the encoded amino acid linkers.

**Figure 4.**
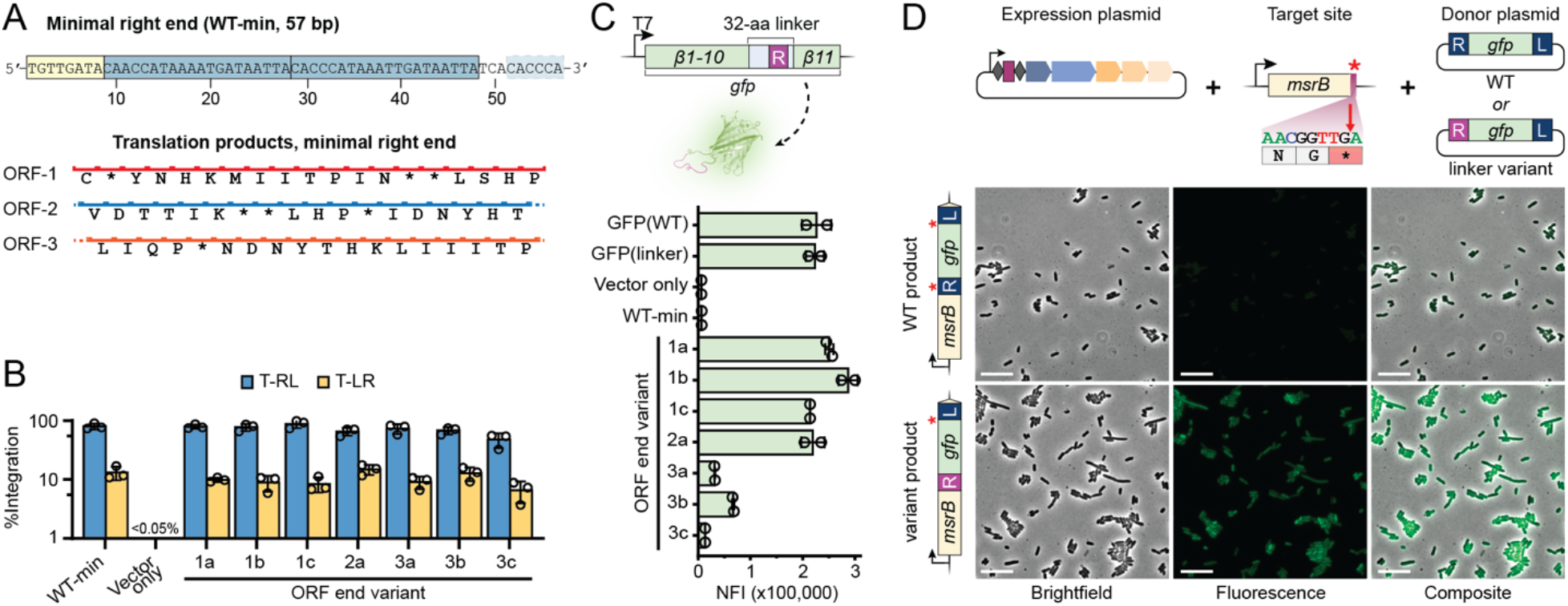
Engineered transposon right ends enable functional in-frame protein tagging. (**A**) An illustration of the minimal transposon right end sequence (“WT-min.”) and the amino acids it encodes in three different reading frames. The 8-bp terminal end (yellow box) and TBSs (blue boxes) are shown. (**B**) Integration efficiencies for individual pDonor variants in which stop codons and codons encoding bulky/charged amino acids were replaced, as determined by qPCR. “Vector only” refers to the negative control condition where pEffector was co-transformed with a vector that did not encode a transposon. (**C**) Select right end linker variants were cloned in between the 10^th^ and 11th β-strands of GFP, in order to identify stable polypeptide linkers that still allow for proper formation and fluorescence activity of GFP. Normalized fluorescence intensity (NFI) was calculated using the optical density of each culture and is plotted for each linker variant alongside wildtype GFP. (**D**) Schematic of a proof-of-concept experiment in which the endogenous *E. coli* gene *msrB* is tagged by targeted, site-specific RNA-guided transposition (top). Fluorescence microscopy images reveal functional tagging of MsrB with the linker variant right end, but not the WT, stop codon-containing right end (bottom). Scale bar represents 10 μm.

GFP naturally consists of eleven β-strands that are connected by small loop regions, and a prior study demonstrated that the loop region between the 10^th^ and 11^th^ β-strand can be extended with novel linker sequences while still allowing for proper folding and fluorescence of the variant GFP protein (34). We cloned selected transposon right end variants into the loop region between β-strand 10 and 11 and measured GFP fluorescence intensity after expression of each construct, which revealed a subset of variants that were fully functional (Figure 4C and Figure S5B). Next, we selected the endogenous *E. coli* gene *msrB* for C-terminal tagging in a proof-of-concept experiment (Figure 4D). After generating a pDonor construct that encodes a right end linker variant with an adjacent, in-frame GFP gene lacking a promoter or start codon, we performed transposition experiments and used Sanger sequencing to verify that integration interrupted the endogenous stop codon while placing the linker and GFP sequence directly in-frame. Finally, proper expression of MsrB-GFP fusion proteins was analyzed by analyzing cells via fluorescence microscopy that received either the WT transposon right end or the linker variant, demonstrating that only the modified right end variant elicited the expected cellular fluorescence (Figure 4D and Figure S5C). Together, these data provide the basis for new genome engineering tools that allow for facile, endogenous gene tagging with single-bp control.

### Integration Host Factor (IHF) binds the left transposon end to stimulate transposition

Closer inspection of the transposon left end mutational data revealed a sequence between the two terminal TnsB binding sites (TBSs) that, when mutated, led to reproducible transposition defects (Figure 5A). We noticed that the corresponding DNA sequence perfectly matched a consensus binding sequence for Integration Host Factor (IHF) (49, 50), a heterodimeric nucleoid-associated protein (NAP) that binds to the consensus sequence 5’-WATCARNNNNTTR-3’ and induces a DNA bend of more than 160° (51). First identified as a host factor for bacteriophage λ integration, IHF is also involved in diverse cellular activities including chromosome replication initiation, transcriptional regulation, and various site-specific recombination pathways (52–54). This observation suggested the intriguing possibility that IHF might also play a role in RNA-guided transposition by CAST systems.

**Figure 5.**
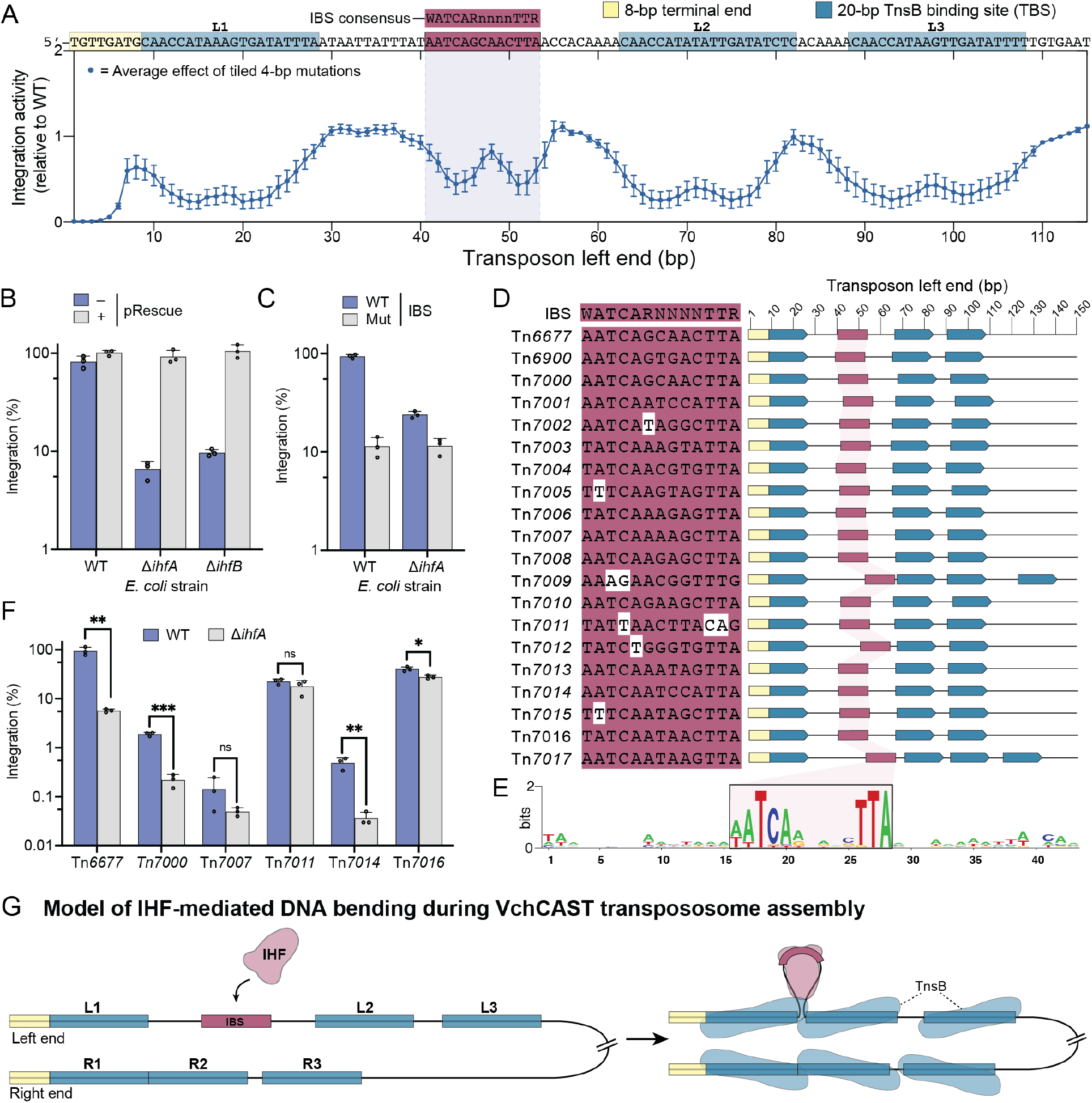
IHF involvement in RNA-guided transposition by VchCAST. (**A**) Library mutagenesis data for the transposon left end. Each point represents the effect of 4-bp mutations, averaged across 4 variants per base. (**B**) Integration activity of VchCAST in WT, Δ*ihfA*, and Δ*ihfB* cells. Integration activity was rescued by a plasmid encoding both *ihfA* and *ihfB* (pRescue). Each point represents integration efficiency measured by qPCR for one independent biological replicate. (**C**) Integration activity when the IHF binding site (IBS) is mutated (Mut), in which all consensus bases within the IBS were modified (from 5’-AATCAGCAAACTTA-3’ to 5’-CCGACTCAACGGC-3’). (**D**) Conservation of the IBS in the transposon left end of twenty Type I-F CAST systems [first described in Klompe et al., 2022] (**E**) Sequence logo generated by aligning the left end sequence of all homologs around the conserved IHF binding site. (**F**) Integration activity in WT and ΔIHF cells for five highly active Type I-F CAST systems. Asterisks indicate the degree of statistical significance:* p ≤ 0.05, ** p ≤ 0.01, ***p ≤ 0.001. (**G**) Model: IHF binds the left end to resolve the spacing between the first two TBSs, bringing together TnsB protomers to form an active transpososome.

To test whether the IHF binding site (IBS) in the left transposon end functions to promote transposition, we first generated IHF knockout strains by mutating either *ihfA* and *ihfB*, and then measured integration efficiency with WT VchCAST. Deletion of either *ihfA* or *ihfB* decreased integration efficiency in the mutant strains by ∼20-fold (Figure 5B), and this effect was completely rescued when we introduced a plasmid encoding recombinant *ihfA* or *ihfB*, confirming the IHF knockouts as causative genetic perturbations (Figure 5B). Interestingly, the reduction in integration efficiency was sensitive to vector design and expression conditions, as integration was less dependent on IHF when the donor DNA was encoded on a separate plasmid from the transposition machinery compared to when the donor DNA was encoded on the same plasmid as the transposition machinery (Figure S6A). When we selectively mutated the conserved IBS residues of a transposon donor, we found that transposition with the mutant left end decreased integration efficiency in WT cells, but not ΔIHF cells (Figure 5C). These experiments indicate that the IBS within the left transposon end is bound by IHF, and that IHF plays a role in stimulating RNA-guided transposition.

We next wondered whether the IHF requirement was conserved across diverse I-F CAST systems, taking advantage of the twenty homologous systems that we recently described (18). Visual examination of the transposon left ends revealed a highly conserved IBS across all homologs (Figure 5D,E), and aligning the sequence between the first two TBSs using Clustal Omega also revealed the IBS consensus as a conserved feature (Figure S6B). To test whether IHF stimulated transposition for these systems, we performed experiments in WT and ΔIHF cells for five other systems and found that only two (Tn*7000* and Tn*7014*) showed a strong IHF dependence (Figure 5F). These data suggest that the IHF dependence may not be conserved across all I-F CAST systems, though the level and length of protein overexpression in our transposition assays likely also affect these results.

Given the involvement of IHF and, more generally, the importance of donor/target DNA supercoiling and topology for other mobile elements (55, 56), we decided to broadly investigate whether other *E. coli* NAPs might play a role in transposition. After generating individual knockouts of 5 additional NAP genes (*ycbG, hupA, hupB, hns*, and *fis*) and measuring integration efficiency within these mutant backgrounds, we found that only the loss of *fis* affected transposition, decreasing integration efficiency by 2-fold (Figure S6F). When we tested the same cohort of NAP knockouts for transposition with the prototypic Tn7 system, IHF had no effect whereas Fis again influenced integration efficiency, though with a ∼4-fold increase in the knockout strain (Figure S7B). Fis (factor for inversion stimulation) plays diverse roles in altering DNA topology, mediating DNA inversions, and regulating gene expression (57–59); these varied roles, and the lack of a clearly defined consensus sequence, make it difficult to know how Fis impacts transposition in either system, or whether changes in integration efficiency might instead be indirect effects. Interestingly, our amplicon-sequencing detection approach for *E. coli* Tn*7* transposition also yielded new information about the nature of DNA integration products for the well-studied TnsABCD pathway. Whereas prior studies concluded that TnsD binding defines a single integration site downstream of the essential *glmS* gene (60–62), we observed surprisingly heterogeneous insertion patterns that sampled a wider sequence space, including rare but reproducible transposition products in the less-common T-LR orientation (Figure S7C). These findings highlight the value of deep sequencing to thoroughly and unbiasedly query the range of potential integration products for a given transposable element.

Lastly, we decided to investigate whether IHF might also bias the orientation of transposon integration for CAST systems, since the IHF binding site (IBS) is uniquely present within the transposon left end. After testing bidirectional transposition for two CAST systems in both a WT and ΔIHF strain of *E. coli*, we found that although the loss of IHF did not affect orientation preference for VchCAST, its loss reversed the dominant orientation for Tn*7000* from T-RL to T-LR (Figure S6C). This result raises the intriguing possibility that IHF may be involved in establishing a transpososome architecture that controls the directionality of DNA insertions, at least for some systems. Previous work with the prototypic Tn*7* system found that transposon substrates with two right ends were competent for integration whereas two left ends were not (13), and we wondered whether a symmetric VchCAST donor with two right ends would similarly be competent for transposition while also eliminating IHF dependency. In agreement with this hypothesis, the loss of IHF had no impact on transposition with a substrate containing two transposon right ends, which was integrated without orientation bias, while a substrate containing two left ends exhibited severely reduced integration efficiency that retained a dependence on IHF (Figure S6D,E). Overall, our data support a model (Figure 5G) in which IHF binds the region between TBSs L1 and L2 to bend the transposon left end and drive DNA integration, akin to the proposed role of HU in Mu transposition [12].

## DISCUSSION

RNA-guided DNA integration by CRISPR-associated transposons depends on diverse, sequence-specific nucleic acid determinants. Focusing on VchCAST, a highly efficient and accurate CAST system derived from *Vibrio cholerae* (also known as Tn*6677*) (24, 31), we employed high throughput screening methods to systematically investigate and characterize these sequence requirements in this study. We first determined the minimal transposon sequences needed for robust activity and validated the importance of each transposase binding site (TBS) found within both left and right ends. Interestingly, our data revealed a broad degree of tolerance to mutagenesis of individual TBSs, a feature corroborated by recent TnsB transposase-DNA structures that show interactions mainly with the DNA backbone rather than specific nucleobases (37–39). The presence of multiple binding sites within each transposon end might allow for accumulative specificity and affinity, and likely play a role in regulating transposition frequency. Our results furthermore suggest that the asymmetric nature of the two transposon ends controls the idiosyncratic preferences of a given element for integrating in one orientation over another.

We uncovered additional regions within the transposon ends that drastically affect integration efficiencies, including a sensitive region within the left end that ultimately revealed a conserved binding site for integration host factor (IHF). Transposition assays with perturbations of the IHF binding site, and in *E. coli* strains lacking IHF, demonstrated that IHF is critical for efficient transposition of VchCAST and some, but not all, homologous Type I-F CAST systems, at least under the expression conditions we tested. Since IHF is known to facilitate extreme bending of the bound DNA (51, 54), we propose that IHF is important for the proper quaternary organization of the transpososome. This hypothesis is further supported by transposon end variants containing alternate spacing between the TBSs, which revealed a conserved periodicity that is consistent with the helical nature of double-stranded DNA. It is striking that, although Type I-F CASTs rely on a multitude of transposon-encoded genes, diverse DNA sequence determinants, and potential additional host-encoded factors, heterologous assays in *E. coli* with twenty CASTs from a range of gammaproteobacteria revealed active transposition for all (18). How and why mobile genetic elements would evolve dependencies on host-specific factors are questions that encourage further research into the regulation of transposition and search for additional accessory factors (63), especially in native host organisms.

We also unbiasedly analyzed sequence biases at the site of integration and found a clear preference for insertions into sites containing a central 5’-YWR-3’ motif, with additional nucleotide preferences 3-bp upstream and downstream of the TSD in regions that appear to make direct contacts with the TnsB transposase from a Type V-K CAST (41). Remarkably, by projecting this new information onto the integration site patterns we previously obtained for a panel of genomic target sites in *E. coli*, we were able to explain the observed product heterogeneity, thus enabling guide RNA selection with high predictability for integration products at single-bp resolution. Finally, we exploited our dataset on transposon end mutability and integration site preference to design modified transposon variants that enabled in-frame tagging of endogenous protein-coding genes. In a proof-of-concept experiment, we tagged the endogenous *E. coli msrB* protein with GFP through modification of a short transposon right end and an in-frame *gfp* gene, and similar efforts should enable in-frame tagging in other cell types, where transposon end ‘scars’ are converted into functional sequence modifications.

Collectively, our work demonstrates the power of combining rationally designed libraries with deep sequencing approaches. We reveal new insights on the molecular mechanism of RNA-guided transposition while also building a register, at single-bp resolution, of which bases can and cannot be mutated for engineering purposes. These new insights inform future studies of both the biology and application potential of CAST systems.

## AVAILABILITY

High-throughput sequencing data are available at the National Center for Biotechnology Information (NCBI) Sequence Read Archive (BioProject Accession: PRJNA919078). Custom scripts used for analyses of high-throughput sequencing data are available at GitHub (https://github.com/sternberglab/Walker_Klompe_etal_2023). Datasets generated and analyzed in the current study are available from the corresponding authors on reasonable request.

## SUPPLEMENTARY DATA

Supplementary Data are available at NAR online.

## ACKNOWLEDGEMENT

We thank S.R. Pesari for laboratory support; N.E. Sanjana for helpful discussions about pooled library experiments; M.A. Hydorn and J.E. Dworkin for fluorescence microscopy support and microscope access; L.F. Landweber for qPCR instrument access; and the staff at the JP Sulzberger Columbia Genome Center for NGS support.

## SUPPLEMENTARY FIGURES

**Figure S1.**
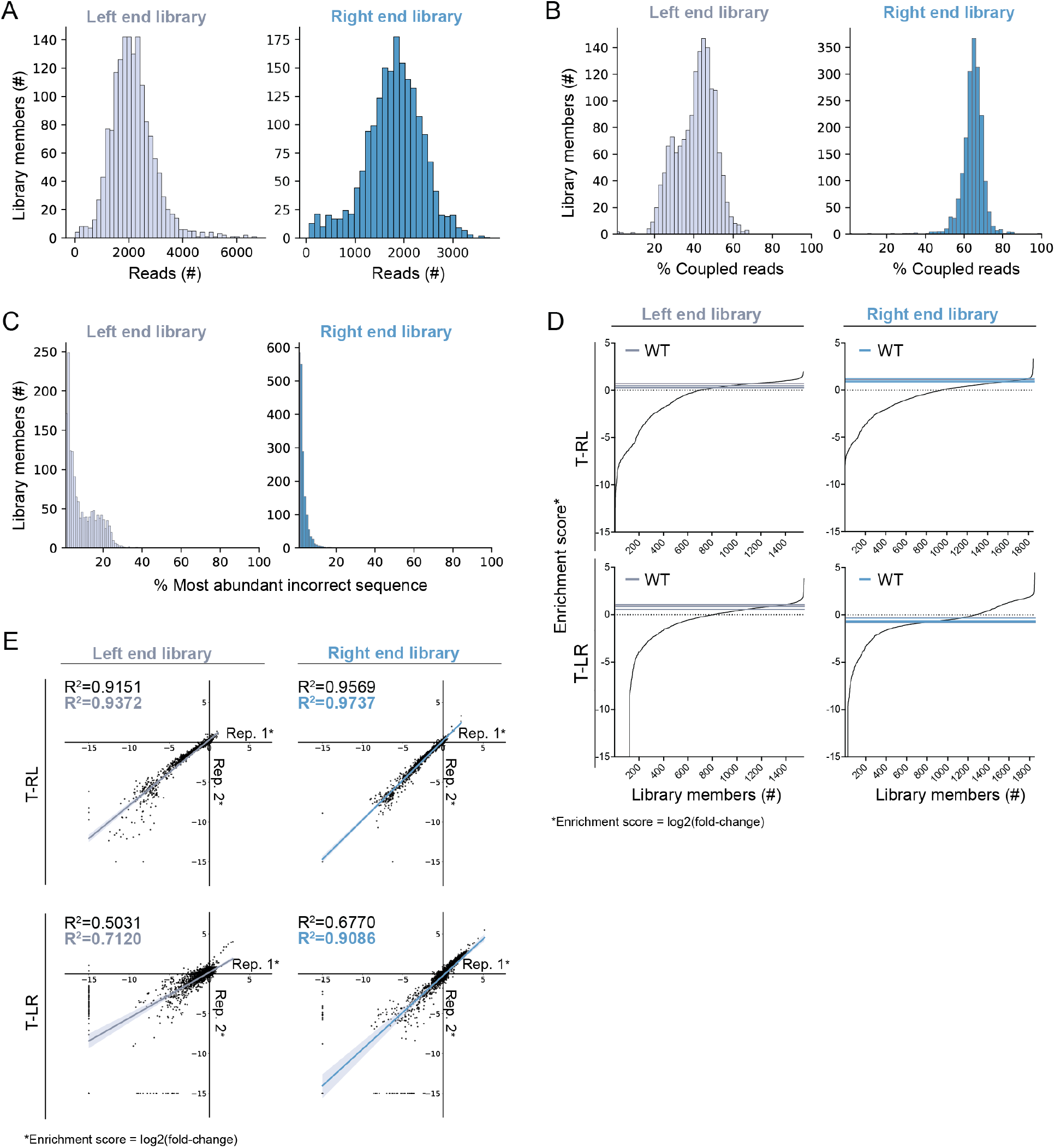
Sequencing and characterization of pDonor right end and left end pooled libraries. (**A**) Histogram showing read counts for each of the input libraries, as defined by barcode sequences. All library members are represented in both the transposon left end and right end libraries. (**B**) Histogram showing the percentage of each library member’s high-quality reads in which the correct barcode is coupled to the correct transposon end sequence. Library members are identified by their barcodes. (**C**) Histogram showing the highest percentage of each library member’s uncoupled reads mapping to a single incorrect sequence. In other words, for a given library member, we selected the incorrect (uncoupled) sequence with the highest read count and expressed that read count as the percent of total reads for that library member. These analyses demonstrate that only a small minority of all barcode reads for a given library member are associated with an incorrect (uncoupled) transposon end sequence. (**D**) All enrichment scores for library members in either integration orientation, for both the left end and right end libraries. Enrichment scores were calculated by dividing the abundance of each member in the output library by its abundance in the input library, and then taking the log2 transformation of that value. Library member dropouts were arbitrarily assigned a score of -15, which fell below the minimum enrichment score across all samples, in order to be plotted on the same graphs. (**E**) Correlation between two independent biological replicates for the transposon left and right end library transposition experiments. For each graph, the upper R2 value (black) includes enrichment scores for all transposon end variants, where dropouts were arbitrarily set to -15. The lower R2 value (colored) includes only the enrichment scores for transposon end variants that were detected in both output libraries.

**Figure S2.**
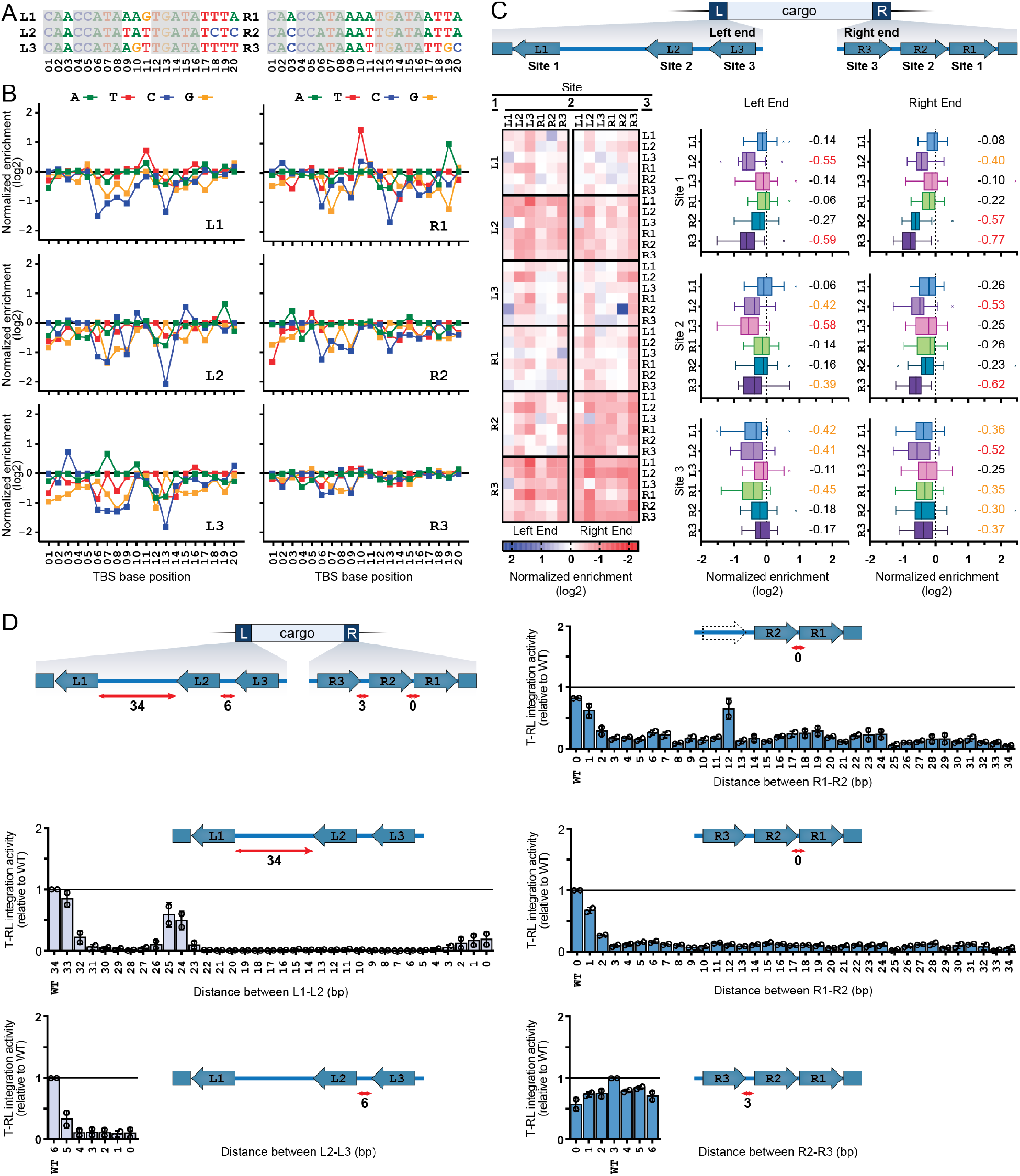
Sequence and spatial requirements of VchCAST TBSs. (**A**) Sequence conservation among the six bioinformatically predicted TBS sequences, with nucleotides conserved among all six sites highlighted in gray. (**B**) Integration activity for mutagenized TBS sequences at individual binding sites, shown as the mean of two biological replicates. Integration activity is represented as the library variant enrichment score normalized to WT. (**C**) Schematic representation of the transposon end architecture (top). Enrichment of individual transposon end variants for which the TBS were shuffled are shown as a heatmap (left). The overall effect of each TBS is represented in a boxplot for the individual sites within both the left and right transposon ends, including their numerical mean (right). (**D**) Schematic representation of the spacing in between the TBS sequences of the transposon left and right ends (top left). Integration efficiencies, calculated from enrichments within the larger transposon end library dataset, are shown for alternative spacing between the TBS sequences of the left and right end sequences.

**Figure S3.**
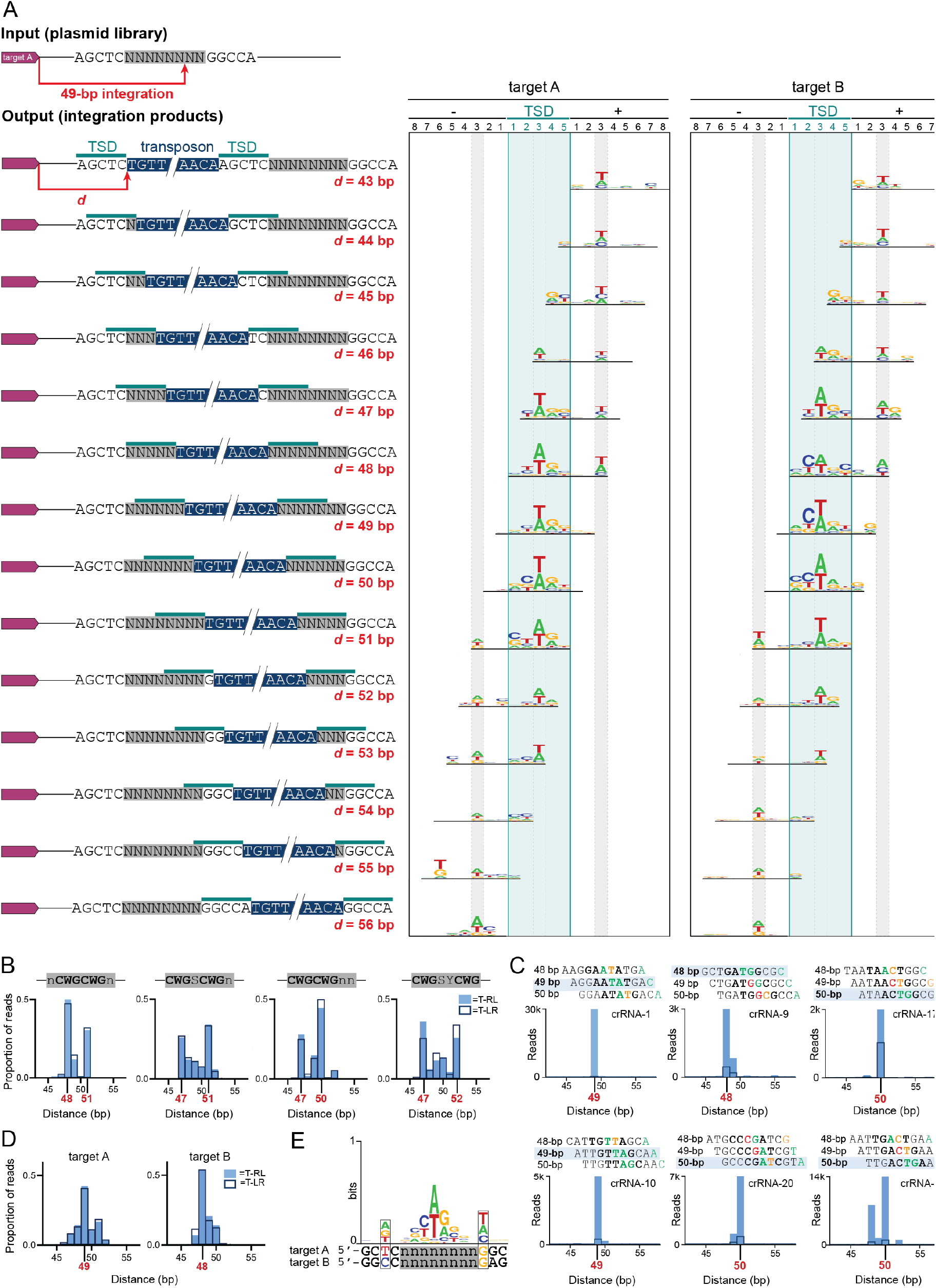
Transposase sequence preferences at the site of DNA integration. (**A**) Schematic of target A integration products, with corresponding sequence logos of enriched sequences at each integration position. Sequence logos were generated by selecting all sequences with 4-fold enrichment in the integrated products compared to the input libraries. The y-axis of each sequence logo was set to a maximum of 1 bit. (**B**) Integration site distance distribution for degenerate sequences containing multiple preferred CWG motifs, with preferred distances indicated in red. (**C**) Integration site distance distributions of previously tested genomic target sites, as determined through deep sequencing. The TSD sequence +/- 3-bp is shown for distances of 48, 49, and 50 bp. Integration occurs primarily 49-bp downstream of the target site but can be biased to occur 48- and/or 50-bp downstream due to sequence preferences at the site of integration. The TSD is bold, and favored (green) or disfavored (orange and red) nucleotides according to the preference sequence logo are indicated. (**D**) Integration site distance distribution for two targets, A and B, with preferred distances indicated in red. (**E**) Nucleotide preferences surrounding the degenerate sequence may be responsible for differences in the overall integration site distance distribution.

**Figure S4.**
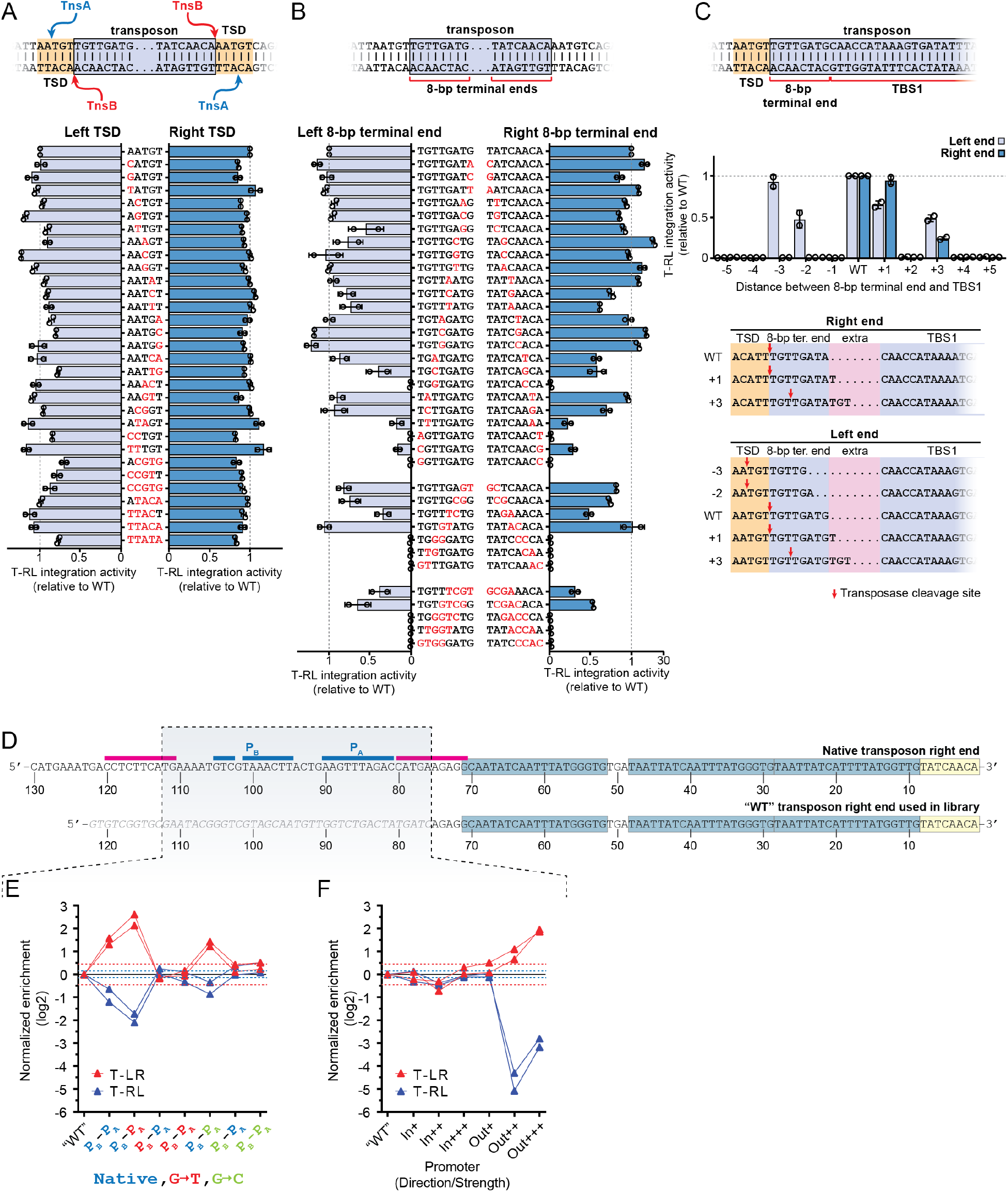
Effect of target-transposon boundary sequences and internal sequences on DNA integration. (**A**) Schematic representation of DNA cleavage by TnsA and TnsB, leading to full excision of the transposon from the donor site (top). Different transposon-flanking sequences were tested on both the left and right transposon boundaries, and integration efficiencies were determined by calculating the enrichment of each library member from within the larger transposon end pool (bottom). (**B**) Illustration of the imperfect 8-bp terminal end sequences for VchCAST (top). Calculated integration efficiencies are plotted for transposon end variants in which either the left or right terminal end sequence was mutated. (**C**) Illustration of the transposon end sequences including the target site duplication (TSD), 8-bp terminal end, and first transposase binding site (TBS1). The specific sequence shown is derived from the VchCAST left end (top). Integration efficiencies relative to WT are shown for transposon end variants in which the distance between the 8-bp terminal end and TBS1 was altered for either the transposon left or right end (middle). Analysis of deep sequencing data revealed TnsB cleavage sites for the right end and left end variants that were functional for transposition; cleavage sites are indicated with red arrows. (**D**) An illustration of WT and modified transposon right end sequences. The 8-bp terminal end (yellow boxes), transposase binding sites (blue boxes), and palindromic sequences (blue and pink lines), are indicated. The native sequence encompasses 130 bp from *V. cholerae* Tn*6677*, whereas only 75 bp were used in the “WT” sequence used in library experiments. (**E**) Integration activity of right end library variants, in which the palindromic sequence was altered. Integration activity is represented as the library variant enrichment score normalized to WT. Each variant included a distinct combination of palindromic sequences P_B_ and P_A,,_ with the ordering as shown. Blue text (“native”) indicates the native palindromic sequence. Orange text (“G-T”) refers to variants in which palindrome nucleotides were mutated from G to T and A to C. Green text (“G-C”) refers to variants in which palindrome nucleotides were mutated from G to C and A to T. (**F**) Integration efficiencies of right end variants in which different internal promoter sequences point inwards of the transposon (In) or outwards across the transposon end (Out). Promoter strengths are indicated pJ23114 (+), pJ23111 (++), pJ23119 (+++).

**Figure S5.**
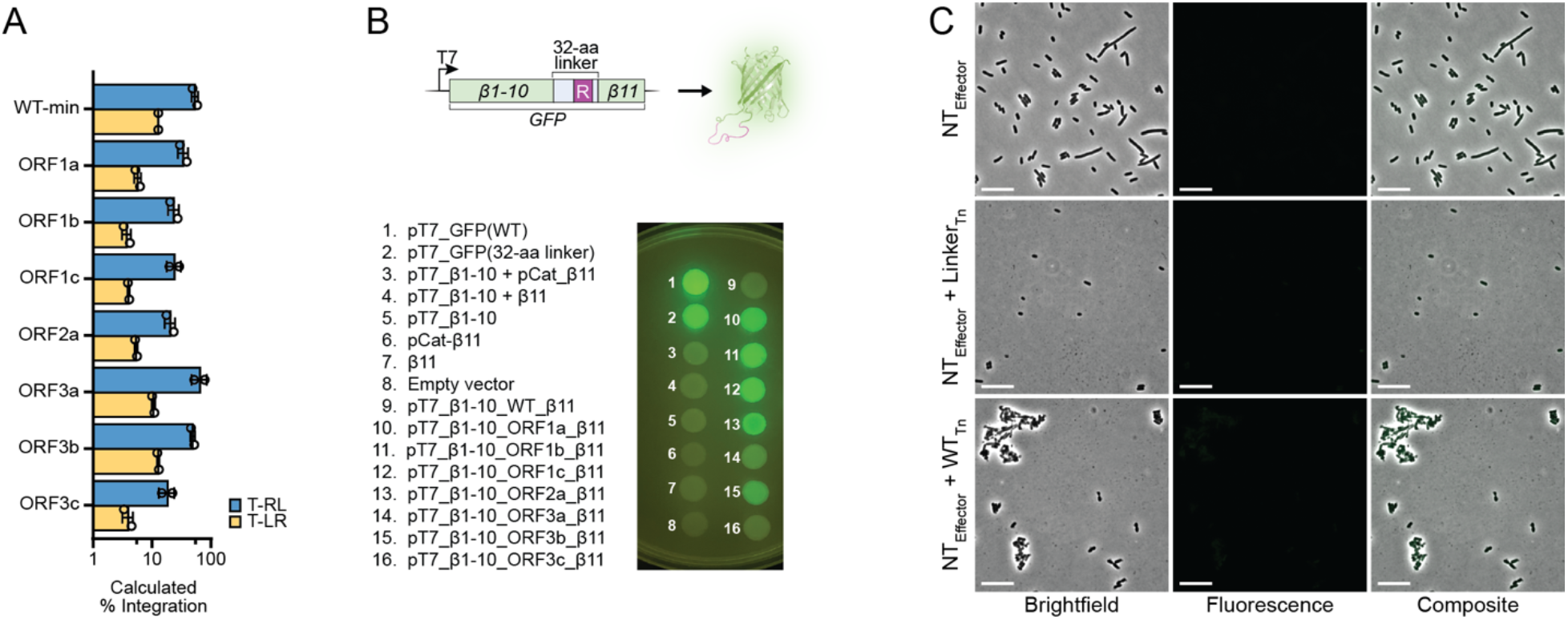
Engineering efforts of the VchCAST right end. (**A**) Integration data for transposon right end variants that were modified to encode functional protein linker sequences in each of three open reading frames (ORF1–3). Integration efficiencies were calculated based on enrichment values within the library dataset. (**B**) Schematic representation of the linker functionality assay in which GFP includes a linker sequence encoded by a mutated right end (top). The fluorescence of *E. coli* cells expressing each of the indicated GFP constructs was visualized upon excitation with blue light (bottom). (**C**) Fluorescence microscopy images of negative control samples for the C-terminal GFP-tagging experiment, showing a brightfield image (left), fluorescence image (center), and composite merge (right). Controls included experiments testing a non-targeting pEffector alone (top) or in combination with either a transposon encoding a functional linker variant (middle) or a wildtype transposon (bottom). Scale bar represents 10 μm.

**Figure S6.**
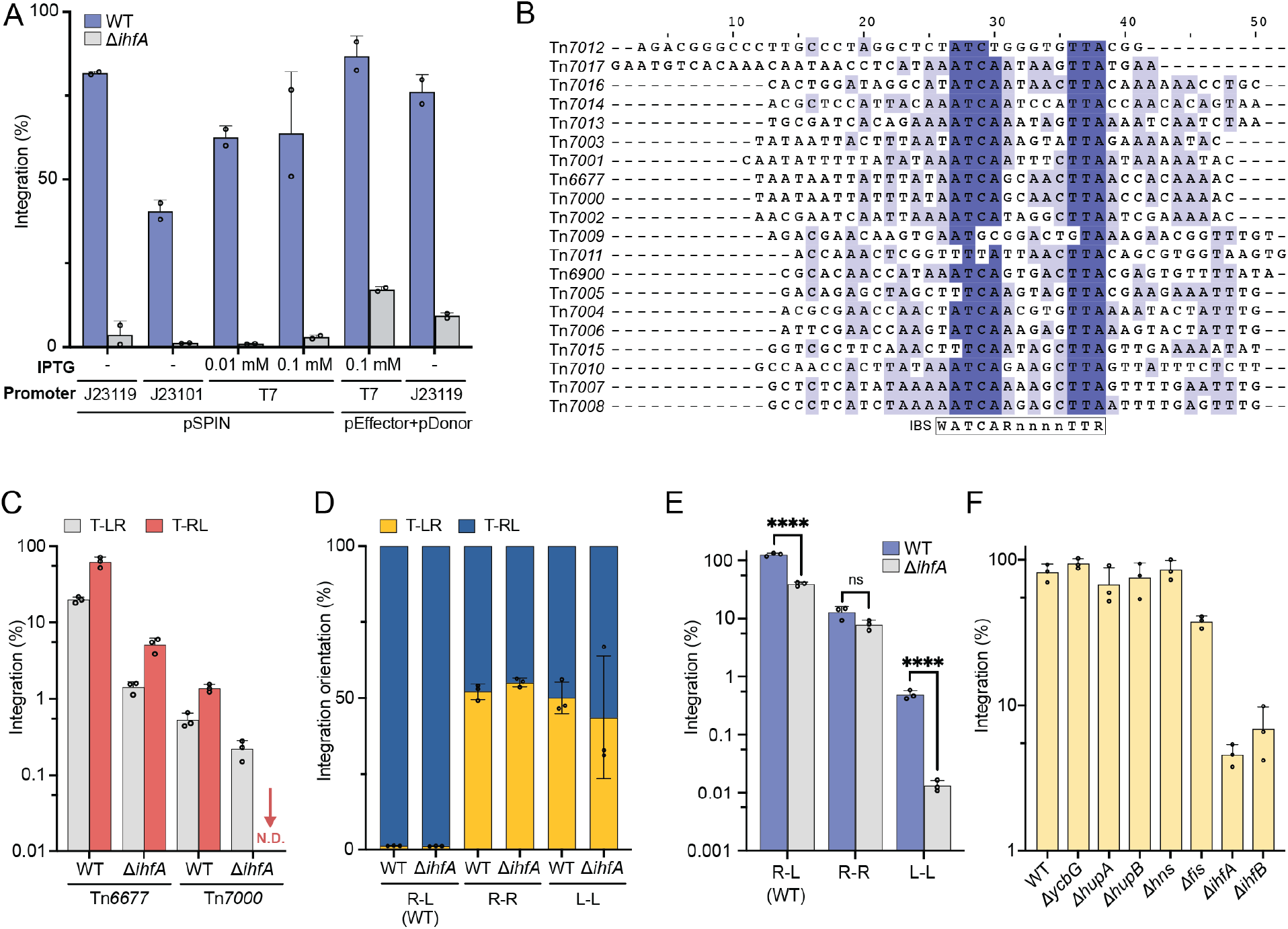
Transposition efficiency of VchCAST and other Type I-F CAST systems in WT and NAP-knockout cells. (**A**) Integration efficiency under different expression systems and induction conditions for VchCAST in WT and Δ*ihfA* cells. pSPIN is a single plasmid that encodes both the donor molecule and transposition machinery, as described in (31). pEffector+pDonor refers to separate plasmids that encode the transposition machinery and donor DNA, respectively. The indicated promoters were also tested, with J23119 and J23101 being constitutively active whereas the T7 promoter is induced by growing cells on IPTG. (**B**) Alignment of the sequence between the first two TnsB binding sites (L1 and L2) in the left end, generated by Clustal Omega and colored in Jalview to highlight conserved residues. The consensus IHF binding site (IBS) is shown below the alignment. (**C**) Integration orientation preference in WT and Δ*ihfA* cells for VchCAST and Tn*7*000. For Tn*7*000, T-RL integration products were not detected (N.D.) after 35 cycles of qPCR, indicating an integration efficiency less than 0.01%. (**D**,**E**) Integration orientation (**D**) and efficiency (**E**) of transposons with symmetric end sequences in WT and Δ*ihfA* cells. R-L refers to a WT-like sequence in which the transposon end identity has not been changed, whereas R-R or L-L refer to transposons in which the left or right end sequence have been mutated to the opposite end sequence, resulting in a transposon with symmetric ends. **F**) Effect of nucleoid associated protein knockouts for VchCAST. Transposition was measured by qPCR after expressing pSPIN in each of the indicated *E. coli* knockout strains.

**Figure S7.**
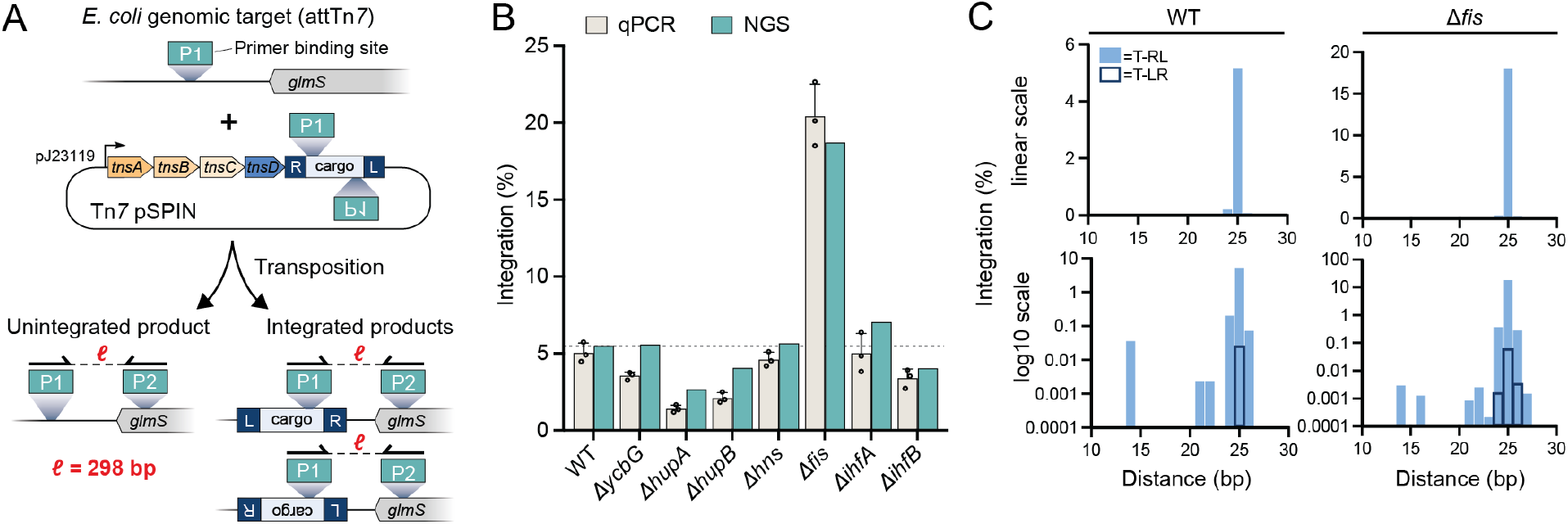
Effect of NAP knockouts on Tn*7* transposition efficiency and fidelity. (**A**) Schematic of NGS-based Tn*7* transposition assay. The transposon cargo encodes genomic primer binding sites (“P1”) adjacent to the right and left ends, such that the NGS amplicon length (“*ℓ*”) is the same for unintegrated products and for integrated products in both orientations. Using this strategy, a single NGS library reports both the integrated and unintegrated products, while avoiding PCR bias that might arise from amplifying products of different lengths or primer binding sites. (**B**) Tn*7* integration efficiencies in the indicated NAP knockout strains are shown, quantified using both qPCR and NGS. The dotted line shows the WT integration value as measured by NGS. Δ*ihfA* or Δ*ihfB* have no effect on integration activity, whereas Δ*fis* increases integration activity ∼4-fold. (**C**) Integration distance and orientation distribution downstream of the *glmS* locus for Tn*7* in WT and Δ*fis* cells. The x-axis refers to the distance in bp between the stop codon of *glmS* and the integration site. For WT and knockout cells, the dominant distance is the canonical 25 bp downstream of *glmS*. The y-axes are shown as linear scale (top) and as log10 scale (bottom), in order to highlight low frequency integration events at non-canonical distances and orientations.

## TABLE LEGENDS

Supplementary Table 1. Plasmids used in this study.

Supplementary Table 2. Oligos used in this study.

Supplementary Table 3. Transposon left end variants and their abundances/enrichments.

Supplementary Table 4. Transposon right end variants and their abundances/enrichments.

